# Finding the right fit: A comprehensive evaluation of short-read and long-read sequencing approaches to maximize the utility of clinical microbiome data

**DOI:** 10.1101/2021.08.31.458285

**Authors:** Jeanette L. Gehrig, Daniel M. Portik, Mark D. Driscoll, Eric Jackson, Shreyasee Chakraborty, Dawn Gratalo, Meredith Ashby, Ricardo Valladares

## Abstract

A longstanding challenge in human microbiome research is achieving the taxonomic and functional resolution needed to generate testable hypotheses about the gut microbiome’s impact on health and disease. More recently, this challenge has extended to a need for in-depth understanding of the pharmacokinetics and pharmacodynamics of clinical microbiome-based interventions. Whole genome metagenomic sequencing provides high taxonomic resolution and information on metagenome functional capacity, but the required deep sequencing is costly. For this reason, short-read sequencing of the bacterial 16S ribosomal RNA (rRNA) gene is the standard for microbiota profiling, despite its poor taxonomic resolution. The recent falling costs and improved fidelity of long-read sequencing warrant an evaluation of this approach for clinical microbiome analysis. We used samples from participants enrolled in a Phase 1b clinical trial of a novel live biotherapeutic product to perform a comparative analysis of short-read and long-read amplicon and metagenomic sequencing approaches to assess their value for generating informative and actionable clinical microbiome data. Comparison of ubiquitous short-read 16S rRNA amplicon profiling to long-read profiling of the 16S-ITS-23S rRNA amplicon showed that only the latter provided strain-level community resolution and insight into novel taxa. Across all methods, overall community taxonomic profiles were comparable and relationships between samples were conserved, highlighting the accuracy of modern microbiome analysis pipelines. All methods identified an active ingredient strain in treated study participants, though detection confidence was higher for long-read methods. Read coverage from both metagenomic methods provided evidence of active ingredient strain replication in some treated participants. Compared to short-read metagenomics, approximately twice the proportion of long reads were assigned functional annotations (63% vs. 34%). Finally, similar bacterial metagenome-assembled genomes (MAGs) were recovered across short-read and long-read metagenomic methods, although MAGs recovered from long reads were more complete. Overall, despite higher costs, long-read microbiome characterization provides added scientific value for clinical microbiome research in the form of higher taxonomic and functional resolution and improved recovery of microbial genomes compared to traditional short-read methodologies.

**Data Summary:** All supporting data, code and protocols have been provided within the article or as supplementary data files. Two supplementary figures and four supplementary tables are available with the online version of this article. Sequencing data are accessible in the National Center for Biotechnology Information (NCBI) database under BioProject accession number PRJNA754443. The R code and additional data files used for analysis and figure generation are accessible in a GitHub repository (https://github.com/jeanette-gehrig/Gehrig_et_al_sequencing_comparison).

**Impact Statement:** Accurate sequencing and analysis are essential for informative microbiome profiling, which is critical for the development of novel microbiome-targeted therapeutics. Recent improvements in long-read sequencing technology provide a promising, but more costly, alternative to ubiquitous short-read sequencing. To our knowledge, a direct comparison of the informational value of short-read and HiFi long-read sequencing approaches has not been reported for clinical microbiome samples. Using samples from participants in a Phase 1b trial of a live biotherapeutic product, we compare microbiome profiles generated from short-read and long-read sequencing for both amplicon-based 16S ribosomal RNA profiling and metagenomic sequencing. Though overall taxonomic profiles were similar across methods, only long-read amplicon sequencing provided strain-level resolution, and long-read metagenomic sequencing resulted in a significantly greater proportion of functionally annotated genes. Detection of a live biotherapeutic active ingredient strain in treated participants was achieved with all methods, and both metagenomic methods provided evidence of active replication of this strain in some participants. Similar taxonomies were recovered through metagenomic assemblies of short and long reads, although assemblies were more complete with long reads. Overall, we show the utility of long-read microbiome sequencing in direct comparison to commonly used short-read methods for clinically relevant microbiome profiling.

## Introduction

The human microbiota consists of trillions of microorganisms colonizing the skin and mucosal surfaces. Much of the diversity and biomass of the microbiota resides in the intestinal lumen, with over 50% of the solid fraction of stool consisting of bacterial cells (1). The human gut microbiota harbors a diverse array of genes, known as the microbiome, that far outnumber human genes and encode a multitude of enzymatic functions lacking in the human genome. The gut microbiome performs essential functions for the human host, including supporting growth and development during infancy and childhood (2), assisting the development and regulation of the immune system (3,4), protecting against pathogens (5), synthesizing vitamins (6), converting indigestible dietary components to usable energy sources (7), and regulating endocrine function and neurologic signaling (8). Perhaps not surprisingly, perturbations to this microbial community are implicated in a range of human diseases, from allergic and autoimmune diseases to cancer and neurologic diseases (9).

Advances in microbiome science have paved the way for a new class of therapeutics that aim to modulate the structure and function of the microbiota to prevent or treat disease. Live biotherapeutic products (LBPs) are live organisms designed to treat, cure, or prevent disease in humans (10). LBPs may contain a single microbe or a consortium of microbes. Unlike traditional therapeutics that exert their effects on a known target receptor or enzyme, LBPs may exert their effects indirectly through impacting the composition and function of the host microbiota (11). Additionally, it has become increasingly clear that the strain is the functional unit of the microbiota, with distinct strains of the same bacterial species frequently displaying unique functional or structural characteristics with important implications for host-microbe interactions (12,13). In this context, strain-level resolution is essential for taxonomic and functional profiling of the microbiota of healthy and diseased individuals to identify candidate drug substance strains as well as for generating high-resolution and actionable data on microbiome dynamics during a microbiome-directed intervention.

Methods to profile the human microbiota have evolved dramatically over the past three decades, with the effort and cost required for DNA sequencing falling precipitously. However, metagenomic sequencing at the depth required to obtain comprehensive taxonomic and functional information from complex microbial communities remains relatively costly. As an economical alternative, amplicon-based 16S ribosomal RNA (16S rRNA) gene sequencing has dominated the microbiome field. Comparison of the 16S rRNA gene sequence allows for an approximation of relatedness among taxa (14). The 16S rRNA gene is approximately 1,500 base pairs (bp) long and contains nine variable regions which can be primed and amplified at lengths compatible with short-read next-generation sequencing (NGS). Limited by read length, the choice of which variable region(s) to sequence has been a longstanding debate due to varying levels of amplification bias (15,16). Overall, short-read 16S rRNA gene sequencing generates community profiles with low taxonomic resolution that are difficult to interpret considering many bacterial genomes harbor multiple polymorphic copies of the 16S rRNA gene (17).

“Third generation” long-read sequencing technology makes it possible to obtain full-length 16S rRNA gene sequences, eliminating bias in choosing a variable region while increasing taxonomic resolution. However, historically high error rates (>10%) per base of long-read sequencing limited its utility in microbiota profiling. Recently, circular consensus sequencing (CCS) has significantly reduced long-read sequencing errors by performing multiple passes of a circularized template molecule (18). CCS of the full-length bacterial 16S rRNA gene significantly improves taxonomic resolution in microbiota profiling, in many cases to the strain level. Sequencing a diverse (>250-member) mock community using CCS sequencing can generate greater than 90% accuracy in species-level classification with accurate measures of relative abundance (19). Long reads are not limited to the 16S rRNA gene; amplicons can be extended to include the internal transcribed spacer (ITS) region and the 23S gene to provide even greater taxonomic resolution (20). Regardless of the platform, 16S rRNA gene sequencing excludes non-bacterial and non-archaeal members of the microbiota, such as fungi, viruses, and eukaryotic cells, and does not provide direct information about the encoded functional capabilities of a microbiome (16).

As the microbiome field has matured, there has been a necessary shift from descriptive observational studies to the mechanistic studies required to support the development of microbiome-targeted diagnostics and therapeutics (21). The application of metagenomic sequencing for translational applications of microbiome profiling has expanded, partly based on improved detection of microbial species and encoded microbiome function (22). While shallow shotgun sequencing (∼0.5 million reads per sample) has been shown to capture the taxonomic and functional diversity of microbiome samples at a fraction of the cost, this approach does not provide sufficient coverage for *de novo* assembly of novel genes and genomes (23). *De novo* assembly is important for the characterization of microbial communities because reference databases contain only a fraction of microbial diversity (24), a fact that has been highlighted by the recent additions of thousands of novel human gut microbiota strains through metagenomic assemblies (25,26). Even with deep short-read shotgun sequencing, *de novo* assembly remains a challenge due to the presence of repetitive regions within genomes, shared genomic regions across different microbes, and the complexity of microbial communities with unknown and uneven representation of both diverse and closely related strains (27). Long-read sequencing can surmount some of the difficulties in metagenomic assembly by spanning repetitive sequences and requiring less assembly; however, reliance on short-read data to overcome high error rates and low coverage has made this approach costly and complex (28). The recent development of CCS to produce HiFi reads (PacBio), which are 99% accurate over 10 kb in length (18), could increase the value and utility of long-read sequencing for microbiome profiling.

The development and regulatory approval of microbiome-targeted therapeutics requires accurate approaches to obtain high resolution taxonomic and functional microbiome data. We applied four distinct sequencing methods to a subset of human microbiome samples collected from participants enrolled in an LBP clinical trial to evaluate their utility for generating actionable data for clinical microbiome applications. Using amplicon-based sequencing, we compare the ubiquitous short-read V3-V4 16S rRNA amplicon sequencing (SRA) to a recently developed long-read 16S-ITS-23S amplicon (LRA) sequencing approach. Using metagenomic profiling, we also compare short-read metagenomics (Illumina, SRM) and long-read metagenomics (PacBio HiFi, LRM) outputs of the same clinical microbiome sample set. As long-read sequencing incurs a greater per-sample sequencing cost, we evaluate sequencing method costs versus value of data obtained about the microbial community, focusing on the two core areas of microbiome sequence data value: taxonomic resolution and functional genomic capacity. We also determine whether introduced therapeutic bacterial strains can be detected during treatment using these distinct approaches. Finally, we compare the genome assemblies from the two metagenomic sequencing methods, assessing the number, completeness, and diversity of metagenome-assembled genomes (MAGs).

## Methods

### Fecal sample collection and processing

After obtaining informed consent, stool samples were collected from participants in Siolta Therapeutics’ Phase 1b clinical trial, STMC-103H-101, investigating the safety and tolerability of the LBP STMC-103H for the prevention of allergic disease. Participants were treated twice daily for 28 days with STMC-103H or placebo, with clinical visits which included fecal sample collection. Study subjects were provided with stool kits with collection containers (BioCollector, The Biocollective). Samples were collected within 24 hours of a scheduled visit, were frozen immediately after collection, and were maintained frozen (≤ −15°C) until processing. To homogenize fecal samples, each sample was weighed and placed in a Seward closure bag (Cat. No. BA6141/CLR), sterile water was added to the sample in a 1:4 ratio, and the bag was placed in a Stomacher 400 Circulator (Seward) and homogenized for 2 minutes at 230 rpm. One-gram aliquots were transferred into labeled cryovials and placed at −80°C prior to DNA extraction.

### SRA (V3-V4 16S rRNA) sequencing and analysis

#### DNA extraction with cetyltrimethylammonium bromide (CTAB)

Fecal sample aliquots (200 mg) were added to ZR BashingBead Lysis Tubes (Zymo Research) with 1 ml CTAB Extraction Buffer (4% hexadecyltrimethylammonium bromide w/v, 100 mM phosphate buffer pH=7.5, 1M NaCl). Sample tubes were loaded into the OMNI Bead Ruptor 24 Elite Bead Mill Homogenizer (OMNI International) and homogenized at 6 m/s for 1 minute and 15 seconds for 4 cycles with 2 minutes dwell time between each cycle. The tubes were centrifuged at 12,000 x *g* for 10 minutes, and 600 μl of the homogenate supernatant was transferred to a sterile 1.5 ml tube. An equal volume (600 µl) phenol:chloroform:isoamyl alcohol (25:24:1, pH = 8) was mixed with each sample followed by a 5 minute rest and centrifugation at 12,000 x *g* for 10 minutes. The aqueous supernatants were transferred to sterile 1.5 ml tubes, and the organic extraction was repeated. After the second centrifugation, 500 µl of aqueous supernatant was transferred to a sterile 1.5 ml tube. 600 µl chloroform was added to the homogenate, and each tube was vortexed and centrifuged at 12,000 x *g* for 10 minutes. About 400 µl of the supernatant was transferred to sterile 1.5 ml tubes, and ⅔ volume of 100% isopropanol was added to each sample, and each sample was vortexed. The tubes were incubated at 4°C for 2 hours to precipitate the DNA. Samples were then centrifuged at 20,000 x *g* for 20 minutes to pellet the DNA. The supernatant was carefully decanted and the pellet in each tube was washed with cold 70% ethanol, mixing with a pipette tip to break up the pellet. Each tube was vortexed briefly, incubated for 5 minutes at room temperature, and centrifuged at 12,000 x *g* for 10 minutes. The ethanol was decanted, and the ethanol wash step was repeated. After the second wash, the ethanol supernatant was decanted and remaining ethanol was removed with a pipette. The tubes with the DNA pellets were left with their caps open in a biosafety cabinet for 15 minutes to allow the remaining ethanol to evaporate. 150 µl TE buffer was then added to each tube, and the samples were vortexed and incubated at 37°C for 30 minutes to solubilize the DNA. DNA concentrations were measured using the Qubit dsDNA Broad Range Assay Kit (ThermoFisher Scientific) on the Qubit 4 Fluorometer (ThermoFisher Scientific), and DNA quality was assessed using the 260/280 and 260/230 ratios measured using a NanoDrop Spectrophotometer.

#### Library preparation and sequencing

DNA samples were normalized to 10 ng/µl in TE buffer. Dual-indexed libraries were prepared following Illumina’s 16S Metagenomic Sequencing Library Preparation Protocol for the Illumina MiSeq System (Part # 15044223 Rev. B). Briefly, the variable V3 and V4 regions of the 16S rRNA gene were amplified using the following primers with Illumina overhang adapter sequences:16S Amplicon PCR Forward Primer = 5’ TCGTCGGCAGCGTCAGATGTGTATAAGAGACAGCCTACGGGNGGCWGCAG 16S Amplicon PCR Reverse Primer = 5’ GTCTCGTGGGCTCGGAGATGTGTATAAGAGACAGGACTACHVGGGTATCTAATCC

PCR amplicons were purified using Omega Mag-Bind Total Pure NGS Beads (Omega Bio-tek, Cat. No. M1378), and each sample was uniquely indexed using Nextera XT Index Kit V2 indexing primers (Illumina, Cat. No. FC-131-2001). Indexed libraries were purified with Omega Mag-Bind Total Pure NGS Beads, and libraries were quantified using the Qubit dsDNA High Sensitivity Kit (ThermoFisher Scientific, Cat. No. Q33231). Amplicon libraries were pooled equally by DNA concentration. The amplicon pooled library was prepared for sequencing with a 5% PhiX spike-in and sequenced on a MiSeq using MiSeq Reagent Kit v3 600 cycle.

#### Data analysis

Demultiplexed fastq files were imported into QIIME 2 for sequence processing and analysis (29). Reads were quality filtered, merged, denoised, and chimeras removed using DADA2 (30) in QIIME 2 (*qiime dada2 denoise-paired*; adjusted parameters: --p-max-ee-f 4, --p-max-ee-r 4, -- p-trim-left-f 17, --p-trim-left-r 21, --p-trunc-len-f 268, --p-trunc-len-r 214). The resulting table contained the number of reads in each sample assigned to amplicon sequence variants (ASVs). ASVs were assigned taxonomy using naive Bayes classifiers trained on Greengenes 13_8 99% OTUs (31) or SILVA version 132 99% OTUs (32) extracted for the target V3-V4 region. ASVs with fewer than 10 reads across all samples were filtered out.

For detecting the active ingredient strain, DSM 33213, from Siolta’s STMC-103H LBP in samples, the number of reads in each sample assigned to DSM 33213’s 408-bp-long V3-V4 ASV were counted and divided by the total number of reads per sample.

### LRA (16S-ITS-23S) sequencing and analysis

#### DNA extraction

To compare the impact of DNA extraction method on microbiome profiles, DNA was extracted from aliquots of each sample using the AllPrep PowerFecal DNA/RNA kit (Qiagen, Cat. No. 80244), the CTAB buffer-based protocol, and the Shoreline Complete DNA extraction as part of the StrainID kit (Shoreline Biome). Each sample was prepared in duplicate using the StrainID kit, and 10 ng of DNA from each sample extracted using the AllPrep PowerFecal kit and CTAB buffer-based DNA extraction were added to the StrainID plate prior to amplification. In addition to fecal samples, both DNA (10 ng) and cells (10^8^ and 10^9^) from STMC-103H’s strains were included in the StrainID library preparation as references.

#### Library preparation

LRA library preparation and analysis were performed as previously described (20) and summarized below.

Approximately 5 mg of fecal material or (10 µl isolate culture) was used as input into the StrainID kit. Fecal sample and isolate culture DNA were isolated per manufacturer’s instructions. Briefly, 50 µl of reconstituted lysis reagent was added to each sample. Subsequently, 50 µl of 0.4 M KOH solution was added and the samples were heated to 95°C for 5 minutes to lyse the cells. The plate was spun briefly to pellet the fecal debris, and 50 µl of the supernatant was transferred to a clean plate. 50 µl of DNA Purification Beads were added, the DNA was allowed to bind, and the pellets were washed with 70% ethanol. DNA was eluted in TE, and 10 µl eluted DNA was transferred to the corresponding well in the PCR plate. At this point, 10 ng of DNA in 10 µl TE from replicates with DNA extracted using other methods was added to separate wells in the PCR plate. 2x PCR mix was added to all wells, and PCR was performed as per instructions to amplify and barcode each sample. Samples were pooled and purified with MinElute spin columns (Qiagen, Cat. No. 28004).

#### DNA sequencing

The amplicon library was created using the SMRTbell Express Template Prep Kit 2.0 (PacBio, Cat. No. 100-938-900) as per manufacturer’s instructions. The library was sequenced on the Sequel II System (PacBio). After sequencing, CCS fastq reads were generated from raw data using default settings for PacBio SMRTlink software.

#### DADA2 installation and analysis

Detailed instructions in the Shoreline Biome ‘Expert User Guide’ for DADA2 is available from https://www.shorelinebiome.com/. Briefly, the following were downloaded and installed on a local computer:

1. SBanalyzer version 3.0.14 for Windows /Centos7/Centos8: https://www.shorelinebiome.com/
2. R version 3.6.1: https://www.r-project.org/
3. R packages DADA2, Biostrings, ShortRead, ggplot2, reshape2, gridExtra, phyloseq, RColorBrewer, biomformat
4. Python 3: https://www.python.org/downloads/

SBanalyzer 3.0 was used to demultiplex and classify CCS fastq reads from the Sequel II sequencing run. SBanalyzer 3.0 includes two StrainID pipelines, StrainID_PacBio_species and PacBio_demux_notrim. The demux_notrim pipeline demultiplexes reads and outputs demultiplexed fastq files with sequences that retain their primers and barcodes. The quality information and intact primer sequences are required by DADA2.

#### Read demultiplexing and taxonomic identification

The original multiplexed CCS fastq file was processed with SBanalyzer 3.0 to demultiplex and classify the reads with the StrainID_PacBio_species pipeline. This pipeline demultiplexes reads using barcode sequence pairs and trims primer and barcode sequences, leaving only the biological sequences. It then maps the trimmed, demultiplexed reads to a local copy of the Athena database and classifies each read. This pipeline outputs several files in minimal cleanup mode, two of which are used in subsequent steps of this procedure: a .taxonomy file associating each read ID with a taxonomy, and a .groups file associating each read ID with a sample.

#### Denoising with DADA2 in R

The untrimmed concatenated fastq files were processed using DADA2 in R. The reads were primer trimmed using removePrimers with max.mismatch=2 and allow.indels=TRUE, and reads lacking recognizable forward or reverse primers were discarded. Trimmed reads were then filtered using the fastqFilter command with minLen=1900, maxLen=3000, maxN=0 and maxEE=2. The trimmed and filtered reads were dereplicated using the derepFastq command, producing a derep-class object containing the unique nucleotide sequences. This dereplicated set of sequences was used to build an error model using the learnErrors command. Dereplicated sequences then went through ASV inference and error correction using the dada command with OMEGA_A=1e-40, OMEGA_C=1e-80, and pool=TRUE, producing a dada-class object containing ASV sequences and their associated frequencies. A sequence table containing the sequence and frequency of each ASV in its associated samples was built from the dada-class object using the makeSequenceTable command. The ASV sequences were finally extracted from the sequence table into fasta format for taxonomic identification with SBanalyzer and the sequence table was converted into rich BIOM format for import into QIIME 2 using the commands make_biom and write_biom.

#### Chimera quality control

The sequence table was used to identify possible chimeric ASVs using the isBimeraDenovo command with minFoldParentOverAbundance=3.5. The total percentage of reads represented by potentially chimeric ASV’s was calculated in R.

#### ASV taxonomic classification

ASV sequences were input in fasta format into SBanalyzer 3.0’s command line sbsearch utility to map them to the Athena database and assign taxonomy. For ASVs which were unclassified at the strain or species level, possibly originating from novel organisms, further analysis was applied to group sequences into putative species or strain classifications. ASVs which were unclassified by SBanalyzer 3.0 were instead classified using SBanalyzer 2.4, and the resulting classifications were appended to the output .taxonomy file.

#### PICRUSt2 analysis for prediction of metagenome functions

Reads were clustered and polished using a reimplementation of the NanoCLUST pipeline (33) to generate the representative sequences and associated per-sample abundances.

The python script picrust2_pipeline.py from the PICRUSt2 distribution was applied (34). Since the 16S-ITS-23S sequences are much longer than the 16S sequences in the PICRUSt2 database, a minimum alignment length threshold of 0.5 was used (default for --min_align is 0.8). There were 65 of 3060 sequences that did not meet that threshold and were filtered out. The flags --stratified and --per_sequence_contrib were used.

### SRM (short-read metagenomics) sequencing and analysis

#### Library preparation and sequencing

DNA was extracted using the CTAB method described in the *DNA extraction with cetyltrimethylammonium bromide (CTAB)* section above. Libraries were prepared based on a 20X scaled-down Nextera XT DNA Library Prep Kit (Illumina, Cat. No. FC-131-1024) protocol (35) with some modifications. Briefly, DNA samples were diluted to 0.2 ng/µl in TE buffer. For Tagmentation, 1 µl of 0.2 ng/µl genomic DNA was added to 2 µl of Tagment DNA Buffer and 1 µl Amplicon Tagment Mix. The Tagmentation reactions were mixed gently by pipetting, spun down, and incubated in a thermocycler for 5 minutes at 55°C followed by a hold at 10°C. Then 1 µl of the Neutralize Tagment Buffer was added to each reaction, and the samples were incubated at room temperature for 5 minutes. For PCR-mediated adapter addition and library amplification, 13.8 µl KAPA HiFi HotStart ReadyMix 2X (KAPA Biosystems, Cat. No. KK2602) and 8.8 µl of paired i7 and i5 Nextera XT Index Kit V2 indexing primers (Illumina, Cat. No. FC-131-2001) were added to each Tagmentation reaction, mixed by pipetting, spun down, and amplified on the thermocycler: 72°C for 3 minutes; 98°C for 5 minutes; 14 cycles of 98°C for 10 seconds, 63°C for 30 seconds, and 72°C for 30 seconds; followed by a final extension of 72°C for 5 minutes. For purification, 22 µl of Omega Mag-Bind Total Pure NGS Beads (Omega Bio-tek, Cat. No. M1378) were added to each reaction and mixed, the samples were incubated at room temperature for 5 minutes and then placed on a magnetic stand (Invitrogen DynaMag-96 Side, Cat. No. 12331D) for 2 minutes, allowing the beads and bound DNA to separate out of the solution. With the samples still on the magnetic stand, the supernatant was removed, and the beads were washed twice with 70% ethanol. After the second wash, all residual ethanol was removed with a pipette and the samples air-dried for 10 minutes in a biosafety cabinet. Samples were then removed from the magnetic stand and the beads were resuspended in 30 µl TE buffer and pipetted up and down to mix. The samples were placed on the magnetic stand for 2 minutes, and then the clear supernatant containing the pure DNA libraries were transferred to a clean PCR plate. Libraries were quantified using the Qubit 1x dsDNA High Sensitivity Kit (ThermoFisher Scientific, Cat. No. Q33231) and pooled equally by DNA concentration. The final pool was sequenced on one lane of a NovaSeq 6000 S4 150 bp PE flow cell (Illumina).

#### Analysis

Nextera adapter sequences were trimmed from demultiplexed paired reads using bbduk (BBTools, sourceforge.net/projects/bbmap/) and reads 105 bp and longer were retained. Reads mapping to the human genome (GRCh38) were removed using bbsplit (BBTools, sourceforge.net/projects/bbmap/). Kraken2 was used to assign taxonomy to reads (36) and Bracken was used to estimate species abundance (37). HUMAnN2 was used for functional annotation, using the UniRef90 protein database (38). Clean reads were mapped to STMC-103H genomes using bbmap (BBTools, sourceforge.net/projects/bbmap/), and the fraction of R1 reads mapping to a genome was counted as the read fraction.

Clean reads were assembled using metaSPAdes in paired-read mode with default settings (39). For metagenome assembly evaluation and MAG identification, the HiFi-MAG-Pipeline was used (https://github.com/PacificBiosciences/pb-metagenomics-tools). A brief overview of this workflow is described in the following section on LRM Sequencing and Analysis. The short-read analysis required several modifications. For each sample, a reads file consisting of the pre-processed, interleaved R1 and R2 files was input for coverage calculations. The command used to map reads to contigs was changed to accommodate short reads: minimap2 -ax sr. The filtering parameters for the maximum number of contigs allowed in a bin was changed from 10 to 500.

### LRM (long-read metagenomics) sequencing and analysis

#### DNA extraction, library preparation, and sequencing

DNA was extracted from aliquots of each sample using the AllPrep PowerFecal DNA/RNA kit (Qiagen, Cat. No. 80244). To ensure a larger DNA fragment size for long-read sequencing, samples were homogenized using the OMNI Bead Ruptor 24 Elite Bead Mill Homogenizer (OMNI International, SKU 19-040E) at 4 m/s for 30 seconds for 2 cycles with 5 minutes dwell time between cycles. DNA fragment lengths were measured on the TapeStation 4200 (Agilent, G2991AA) using the Genomic DNA ScreenTape assay (Agilent).

DNA fragment lengths were approximately 8-10 kb, so DNA did not require shearing prior to library preparation. Sequencing libraries for each sample were created using the SMRTbell Express Template Prep Kit 2.0 (PacBio, Cat. No. 100-938-900) per manufacturer’s instructions. Libraries were sequenced with four samples per run on a Sequel II System (PacBio). One library failed sequencing due to a problem with its library preparation (treatment sample from Participant 6). After sequencing, CCS analyses were run using SMRTLink software v10 to produce HiFi reads for each sample.

#### Analysis

Taxonomic and functional profiling was performed using the PacBio pipeline (Taxonomic-Functional-Profiling-Protein; https://github.com/PacificBiosciences/pb-metagenomics-tools). This pipeline uses DIAMOND (40) to align the HiFi reads to a protein database (e.g., NCBI nr). The resulting alignments are interpreted using MEGAN-LR (42,43), which uses a lowest common ancestor (LCA) algorithm and other long-read settings to assign taxonomic and functional annotations to the HiFi reads. The outputs include absolute and normalized taxonomic read counts using a taxonomy from NCBI or the Genome Taxonomy Database (44,45), and counts for functional annotations based on InterPro2GO, SEED, EC, and eggNOG (42).

Metagenomic assemblies were performed for each HiFi read set using HiCanu (46), implemented in Canu v2.0 (47). To apply metagenomic settings, we used the following options: -pacbio-hifi, genomeSize=100m, maxInputCoverage=1000, batOptions=“-eg 0.0 -sb 0.001 -dg 0 -db 3 -dr 0 -ca 2000 -cp 200”. We note that to apply metagenomics settings for the recently upgraded HiCanu v2.1, the following options should be used instead: -pacbio-hifi, genomeSize=100m maxInputCoverage=1000, batMemory=200. To evaluate the assemblies and identify high-quality MAGs, the HiFi-MAG-Pipeline was used (https://github.com/PacificBiosciences/pb-metagenomics-tools). Briefly, this pipeline aligns HiFi reads to assembly contigs to obtain coverage estimates, which are used with MetaBat2 (48) to perform binning. Resulting bins are evaluated using CheckM (49), and quality thresholds are applied to retain high-quality MAGs (>70% completeness, <10% contamination, <10 contigs). The high-quality MAGs are then analyzed using the Genome Taxonomy Database Toolkit (50), which attempts to identify the closest reference genome and assign taxonomy for each MAG.

### Reference matching for metagenomic contigs

For each sample, the reference genomes of the active ingredient strains from the LBP were mapped to the assembly contigs using minimap2 (51). Contigs belonging to the active ingredient strains were identified by creating scatterplots of the resulting alignment lengths and number of matched bases. In the scatterplot, perfectly matched contigs (to a strain) occurred on a line with a slope (m) of 1 and intercept (b) of 0, indicating the number of matched bases was equal to the alignment length (100% identity).

### Comparing taxonomic profiles of different sequencing methods

For comparison of LRA to other methods, one replicate of each sample with the most reads that went through the entire StrainID protocol (i.e., not the replicates with DNA extracted with other methods) was chosen. The pcoa.plot function from the RAM package (version 1.2.0) in R was used to generate principal coordinates analysis (PCoA) plots of Bray Curtis distances between samples using the taxonomy relative abundance tables at the lowest available taxonomy level for each sequencing method. Pairwise Mantel tests were performed in R using the mantel.rtest from the ade4 package (version 1.7-16) with 9999 permutations.

### Number of unique taxa detected across methods

For SRA, the number of unique taxa detected was determined by counting the number of unique ASVs present in each sample from the filtered ASV table (ASVs were filtered out if they had fewer than 10 reads across all samples). For LRA, the number of unique taxa per sample was determined by counting the taxa present at taxonomy level 8 (strain level) from the Athena database assignments. For SRM, taxa assigned by Kraken2 and normalized by Bracken that were present at greater than 0.04% abundance were counted. This was the threshold below which reads in a mock community were assigned to taxa not present in the mock community. For LRM, read counts were obtained for taxa using MEGAN-LR. The normalized counts of taxonomy assignments to the Genome Taxonomy Database (GTDB) were used.

### Taxonomic resolution

For SRA, reads were considered to have species-level taxonomy if the Greengenes taxonomy classifier (or SILVA taxonomy classifier for SILVA analysis) assigned the ASV to a named species (i.e., not unknown or unclassified). For LRA, the replicate of each sample that underwent the entire StrainID protocol and had the greatest number of reads was chosen to assess taxonomic resolution. Reads assigned to taxonomy level 7 with a specific named species (i.e., not unclassified) were counted as having species-level resolution. Reads assigned to taxonomy level 8 with a specific named strain (i.e., not unclassified) were counted as having strain-level resolution. The denominator was the total number of reads assigned to “Bacteria”.

### Differential abundance

The R package MaAsLin 2 (Microbiome Multivariable Associations with Linear Models) (52) was used to identify differentially abundant bacterial families and differentially abundant functional pathways with time point as a fixed effect and participant as a random effect. For LRM, taxonomy assignments based on the Genome Taxonomy Database (GTDB) were used. Family and pathway abundances were normalized with total sum scaling and log transformed.

### Statistics

Statistical analyses were performed in R (version 4.0.4) and Prism 9 (version 9.1.2), as detailed for each analysis.

## Results

### Summary of sequencing and bioinformatic approaches

A summary of the four sequencing methods compared in this analysis is provided in **Table 1**. The commonly applied 515F and 803R V3-V4 primer pair was chosen for short-read amplicon (SRA) library preparation in this analysis because it was shown to be optimal for profiling bacterial communities (54) and maximizing phylogenetic coverage (55). Libraries were sequenced on a MiSeq with paired-end 300 bp reads. For long-read amplicon sequencing (LRA), the full-length 16S rRNA gene plus the ITS region and part of the 23S gene were amplified using the Complete StrainID Kit (Shoreline Biome) (**Fig. 1A**) and libraries were sequenced on the Sequel II System (PacBio) (**Fig. 1B**). Short-read metagenomic (SRM) libraries were sequenced on the NovaSeq 6000 using one lane of an S4 flow cell to generate paired-end 150 bp reads. Long-read metagenomic (LRM) libraries were sequenced on the Sequel II System.

**Table 1.** Summary of sequencing methods for microbiome profiling.

| Method | Sequencing Type | Sequencing Platform | Read lengths (bp) | Number of reads per sample (M) | Output | Data processing required | Cost per sample |
| --- | --- | --- | --- | --- | --- | --- | --- |
| SRA (V3-V4) | Amplicon | Illumina MiSeq | 250 or 300 | 0.03–0.2 | Taxonomic profiling (genus level) | Low | \$ |
| LRA (16S-ITS-23S) | Amplicon | PacBio Sequel II | ~2,500 | 0.01-0.03 | Taxonomic profiling (strain or species level) | Low | \$ |
| SRM (short-read metagenomics) | Metagenome | Illumina NovaSeq 6000 | 150 | 20-100 | Taxonomic and functional profiling, genome assembly and binning | High | \$\$ |
| LRM (long-read metagenomics) | Metagenome | PacBio Sequel II | 7,000 to >10,000 | 1-2.5 | Taxonomic and functional profiling, genome assembly and binning | Moderate | \$\$\$ |

**Fig. 1.**
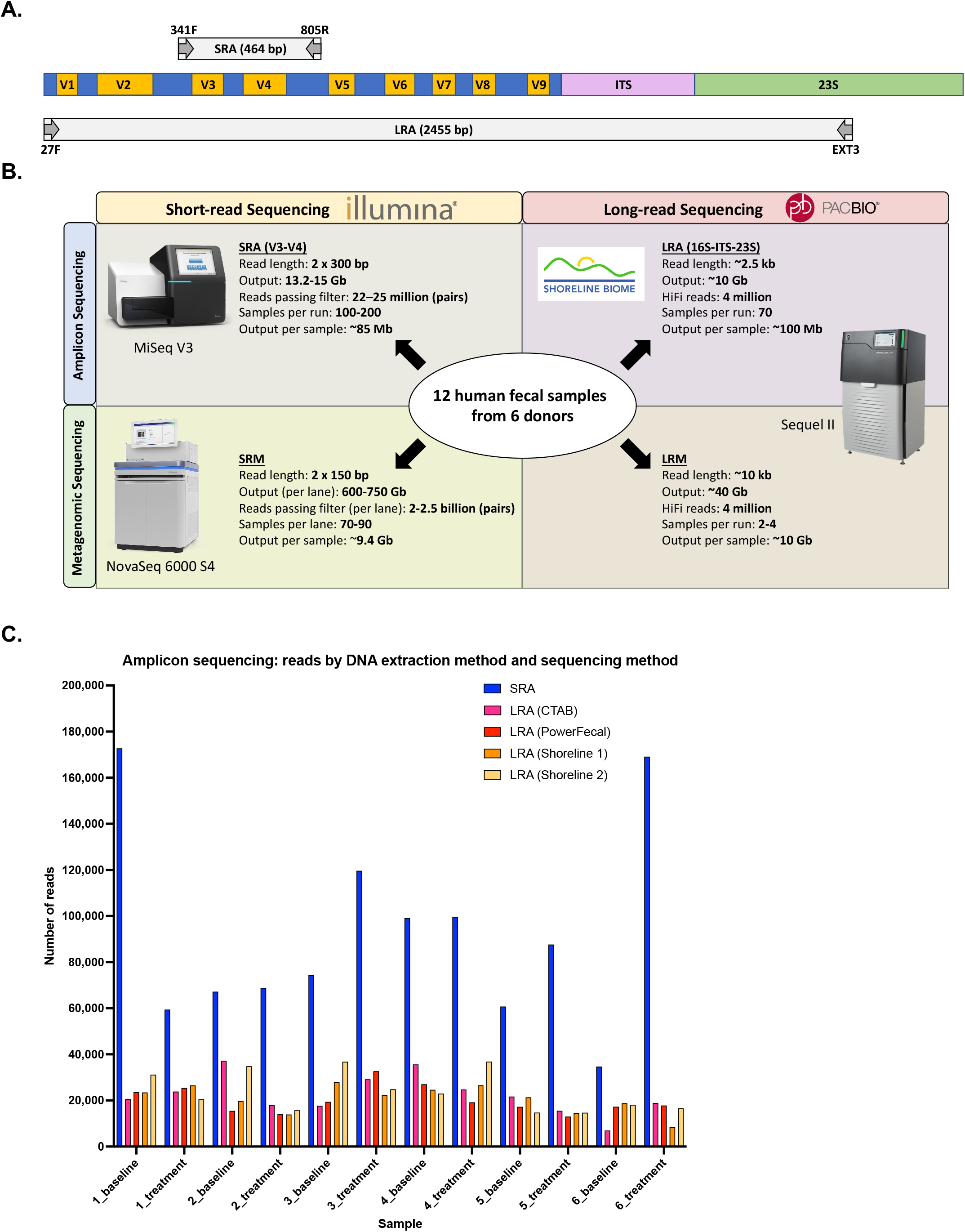
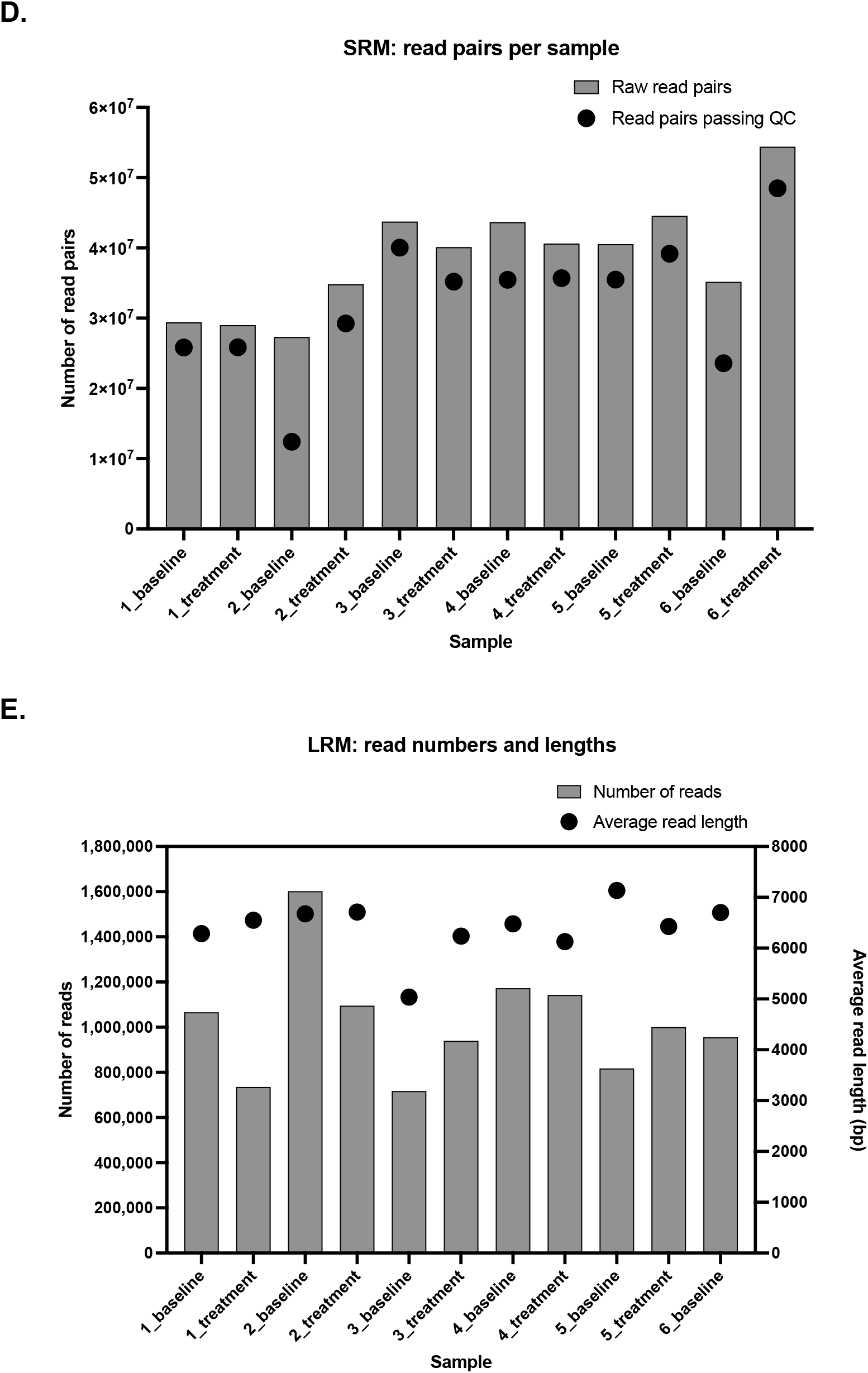
Summary of sequencing methods and output data. **(A)** Comparison SRA and LRA amplicons. SRA consists of a portion of the 16S rRNA gene spanning variable regions 3 and 4, while LRA spans the entire 16S rRNA gene, the internal transcribed spacer (ITS) region, and part of the 23S rRNA gene. **(B)** Summary of the four sequencing methods compared in this analysis, including the sequencing platforms, read lengths, data outputs, and multiplexing capabilities. **(C)** Number of reads passing demultiplexing and filtering for SRA (paired reads are merged during denoising), and the number of HiFi reads passing demultiplexing and classification for LRA by sample replicate. For LRA, the DNA extraction method used for each replicate is denoted in parentheses in the legend. **(D)** The number of raw read pairs and the number of read pairs passing filtering for SRM. **(E)** The number of LRM reads and the average read length per sample.

The DNA sequencing, library preparation, and data analysis differed among the four approaches compared in this analysis (summarized in **Table S1**, detailed in **Methods**). For example, for LRM, DNA must be extracted to generate high-molecular weight DNA from samples. Longer DNA fragments yield longer reads which may contain several full-length bacterial genes, and longer reads improve microbial genome assembly. For short-read sequencing, DNA can be extracted more vigorously to ensure DNA is captured from organisms with tough cell walls, such as fungi and gram-positive bacteria. For data analysis, pipelines are typically optimized for either short or long reads. For example, in this analysis, the assembler metaSPAdes is used to assemble genomes from short metagenomic reads and HiCanu is used to assemble genomes from long metagenomic reads.

Total data output per sample was comparable across the two amplicon sequencing methods and the two metagenomic sequencing methods, with both metagenomic sequencing methods resulting in about 100 times the data of the amplicon sequencing methods (**Fig. 1B**). Notably, the per-sample costs of SRA and LRA are similar, while the per-sample cost of LRM is substantially higher than SRM (**Table 1**). This is due, in part, to the lack of high throughput metagenomic sequencing on the Sequel II System; this experiment included 4 samples per sequencing run (**Fig. 1B**).

### Sample and data summary

The fecal samples analyzed for this comparison were collected from subjects at two timepoints: pre-treatment baseline and 4 weeks after treatment with STMC-103H. Twelve fecal samples from six clinical trial participants were sequenced using four methods: SRA, LRA, SRM, and LRM (**Fig. 1B**). The SRA libraries sequenced on the Illumina MiSeq resulted in an average of 92,766 (± 42,804 SD) combined read pairs per sample after denoising (**Fig. 1C, Methods**). LRA libraries sequenced on the Sequel II resulted in an average of 21,852 (± 7307 SD) HiFi reads per sample (**Fig. 1C**). SRM libraries sequenced on the Illumina NovaSeq 6000 resulted in an average of 38.6 (± 7.8 SD) million 150 bp read pairs per sample; on average, 82.2% (± 13.2% SD) of the raw reads passed filtering criteria (**Fig. 1D, Methods**). LRM libraries sequenced on the Sequel II System produced an average of 1.0 million (± 0.25 SD) reads per sample, with average read lengths per sample ranging from 5 to 7.1 kb (**Fig. 1E**).

### Comparing taxonomic profiles across methods

To determine if short-read and long-read approaches generated comparable taxonomic profiles at higher order taxonomies, all samples were combined and the relative abundances collapsed by phylum and family (**Fig. 2A**). Firmicutes was the dominant phylum across all methods (average abundances of 76%, 54%, 68%, and 81%, for SRA, LRA, SRM, and LRM, respectively) followed by Actinobacteria and Bacteroidetes. The overall proportions of phyla were similar across methods, with some notable differences. Unlike other methods, LRA resulted in a significant proportion of unknown phyla (24%). Most of the reads that mapped to “unknown phyla” (22.8%) were classified at the species level as “bacterium LF-3”. Since bacterium LF-3 is not classified into higher order taxonomies, the reads that mapped to this bacterium remained unclassified at all other levels. SRM resulted in a greater proportion of Actinobacteria compared to the other methods (23% for SRM compared to 8.5% for SRA, 9.3% for LRA, and 8.5% for LRM; one-way ANOVA p<0.0001). Accurate representation of Bifidobacteria depends on thorough mechanical lysis during DNA extraction (56), so the lower representation of Bifidobacteria in the LRA taxonomic profiles could be due in part to the lack of a mechanical lysis step in the Shoreline extraction protocol. Similarly, lower representation of Bifidobacteria in the LRM profiles could be due to the less vigorous bead beating during DNA extraction compared to the DNA extraction for short-read methods. 16S rRNA analysis based on the V3-V4 region and other regions has also been shown to underestimate the abundance of Bifidobacteria (57). SRA and SRM identified a significantly greater proportion of Verrucomicrobia (3.9% for SRA and 0.94% for SRM compared to 0.16% for LRA and 0.41% for LRM; one-way ANOVA p=0.0072). The Verrucomicrobia genus Akkermansia has been shown to be overrepresented in 16S rRNA analysis with the V3-V4 primers (57). SRA and SRM also identified the Archaea phylum Euryarchaeota (1.2% for SRA and 1.3% for SRM), which was not identified with LRA or LRM. However, this phylum was detected when LRM data was annotated using the NCBI database rather than the Genome Taxonomy Database (GTDB). Comparing relative abundances at the lower taxonomic level of family shows broad similarities and some clear differences. Lachnospiraceae was the dominant family across all methods except for LRA, which had a greater proportion of unknown families (Lachnospiraceae abundance 39% for SRA, 4.3% for LRA, 46% for SRM, and 49% for LRM). It is possible that the same bacteria in samples that mapped to the unclassified bacterium LF-3 with LRA mapped to the Lachnospiraceae family with other methods. LRA also had a significantly greater proportion of the family Peptostreptococcaceae (12% for LRA compared to 0.33% for SRA, 1.5% for SRM, and 0.44% for LRM; one-way ANOVA p<0.0001). The families Ruminococcaceae, Bifidobacteriaceae, and Bacteroidaceae, which are abundant in the healthy human gut, were prominent across methods. Both amplicon methods resulted in a higher relative abundance of the family Erysipelotrichaceae (6.3% for SRA and 15% for LRA compared to 2.6% for SRM and 2.0% for LRM; one-way ANOVA p=0.0012).

**Fig. 2.**
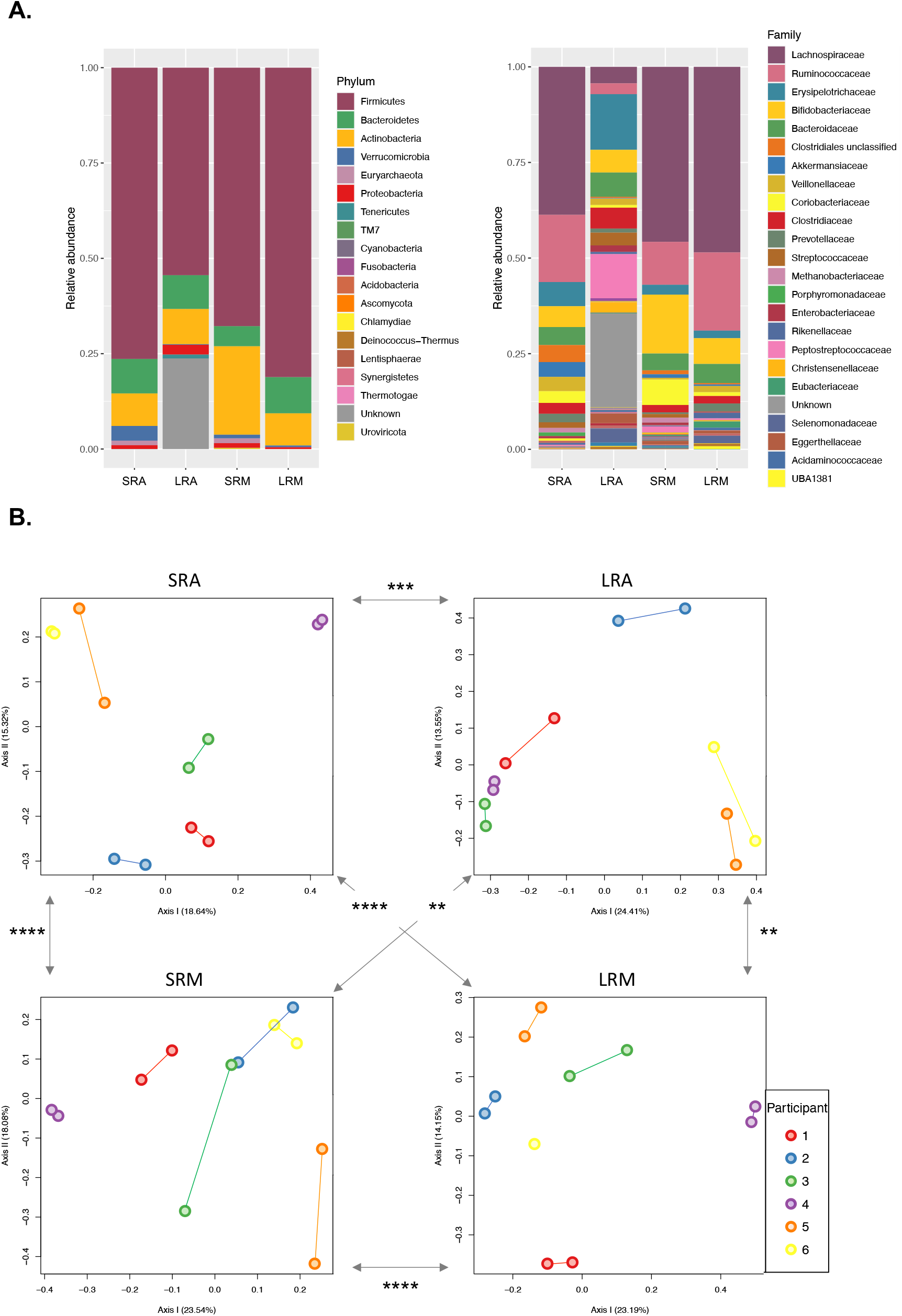
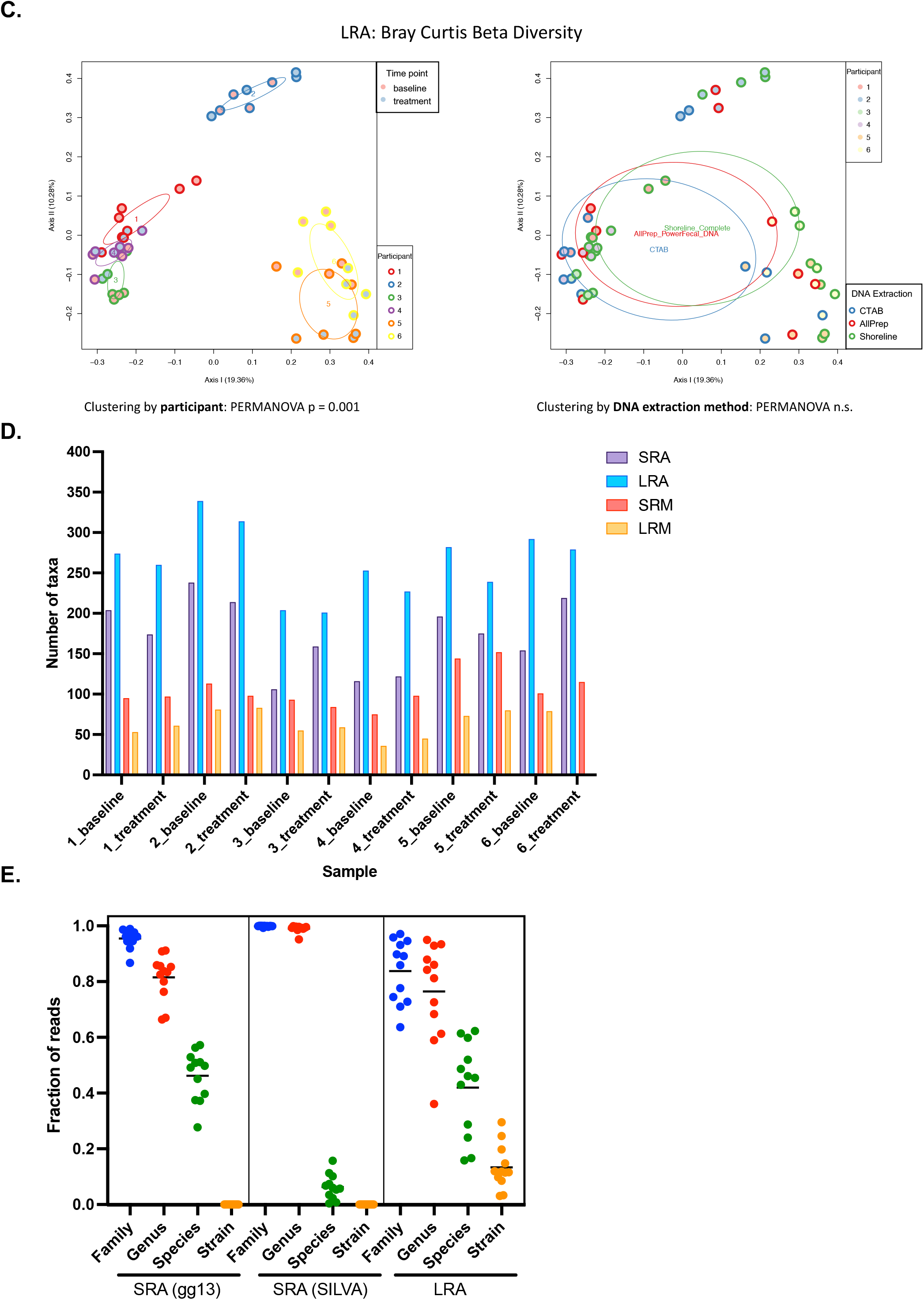
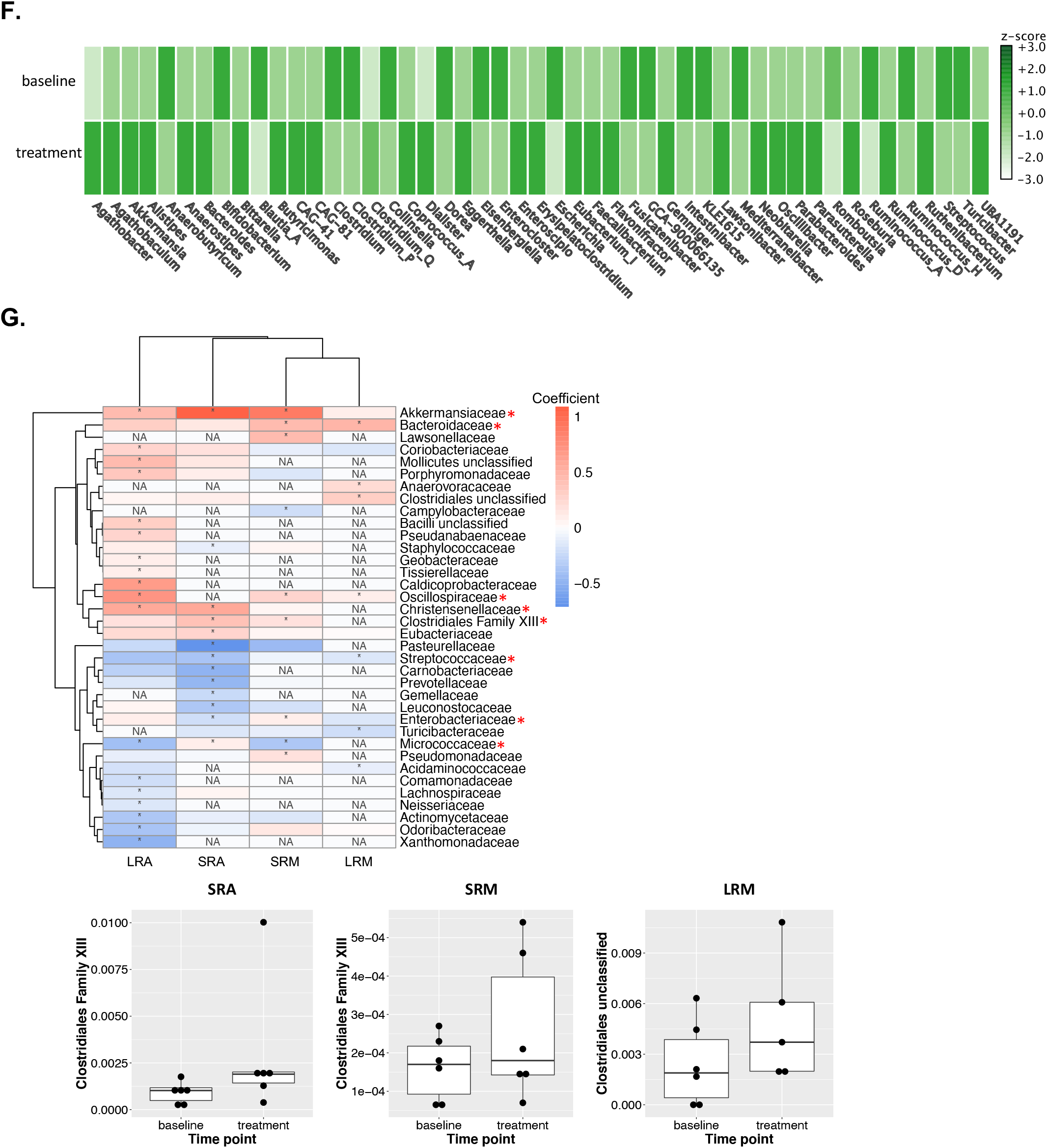
Taxonomic comparisons across methods. **(A)** Taxonomic comparisons across methods at the phylum level (left) and the family level (right). Bar graphs show the relative abundance of each phylum or family across all samples. For the LRA analysis, all four replicates of each sample were included. All phyla and the 24 most abundant families across all methods are shown. **(B)** Bray-Curtis distances between samples were calculated using the taxonomic profiles at the lowest taxonomic level available for each method (species, strain, ASV) and shown with PCoA. Samples are colored by participant, and samples from the same participant are connected by a straight line. Pairwise Mantel tests were performed between Bray-Curtis distance matrices, and significance based on simulated p-values for each comparison are shown with arrows indicating the two distance matrices compared (\**P* < 0.05; \*\**P* < 0.01; \*\*\**P* < 0.001; \*\*\*\**P* < 0.0001). For the LRA distance matrix, one representative replicate was chosen for each sample. Bray-Curtis distances were calculated for all four LRA replicates of each sample that underwent different DNA extraction procedures (CTAB, AllPrep PowerFecal kit, or the Shoreline Complete DNA extraction). Distances between samples are shown on PCoA plots. On the plot on the left, samples are colored by participant and by sample time point, with ellipses for samples from the same participant. There was significant clustering by participant. In the plot on the right, samples are colored by DNA extraction method and by participant, with ellipses for DNA extraction method. There was no significant clustering by DNA extraction method. **(D)** The number of unique taxa detected for each sample, including unknown taxa, at the lowest level of taxonomy for each method. **(E)** The fraction of reads for each sample assigned to the specified taxonomy level for SRA and LRA. Horizontal lines show the mean fraction of reads assigned to each taxonomy level for each method. **(F)** Comparison of levels of genera in Participant 3’s baseline and treatment samples. The z-score shown was computed by MEGAN-LR, where the z-score z(s, t) for any pair of sample (s) and taxon (t) is the z-score (x – μ/s) of t(s) calculated using the list of all values t(s). **(G)** The heatmap shows the families that were significantly differentially abundant between baseline and treatment samples in at least one method (asterisks in heat map denote p < 0.2 for SRA, LRA, and SRM and p < 0.25 for LRM). The red asterisks indicate families that were differentially abundant in more than one method. The relative abundances of the Clostridiales Family XIII/Clostridia unclassified family are plotted in boxplots below the heatmap, showing the three methods in which the family was differentially abundant.

To compare sample taxonomic profiles across methods using the highest taxonomic resolution available to each method, we calculated Bray-Curtis distances (which factors presence/absence and relative abundance of taxa) between samples at the lowest taxonomic level for each method [species, strain, or amplicon sequence variant (ASV)], and compared the distances between samples across methods. Notably, principal coordinates analysis (PCoA) plots of Bray-Curtis distances show that independent of the sequencing method, samples cluster by individual (**Fig. 2B**). Pairwise Mantel tests were performed between Bray-Curtis distance matrices from the four methods and all matrices were significantly positively correlated, suggesting that all four methods preserved relationships between the samples (**Fig. 2B**). Across LRA replicates, samples significantly clustered by participant (PERMANOVA p = 0.001) but not by DNA extraction method (PERMANOVA p = 0.2) (**Fig. 2C**), showing that the DNA extraction methods used did not drastically impact the overall taxonomic profiles.

### Taxonomic resolution across methods

Ideally, microbiota sequencing and analysis should distinguish different strains as unique taxa while minimizing artifacts from DNA library preparation and sequencing. Amplicon approaches detected more unique taxa per sample than metagenomics (**Fig. 2D**). LRA analysis identified the most unique taxa per sample, with an average of 263.7 (± 41.8 SD), while SRA analysis identified an average of 173.1 (± 43.1 SD) unique taxa per sample. For metagenomics, 105.4 (± 22.7 SD) unique taxa at the species level were detected per sample for SRM and 64.1 (± 16.1 SD) for LRM. Within microbiome sequencing methods, there were no significant correlations between the number of reads and the number of unique taxa detected, suggesting that sequencing depth was sufficient to capture community diversity with all methods analyzed (**Fig. S1**).

There was no significant difference between the percent of reads assigned to species for SRA with Greengenes taxonomic assignment (46.2% ± 8.9% SD) and LRA (42.0% ± 16.8% SD) (**Fig. 2E**). However, the percent of species-level assignments was significantly greater with LRA compared to SRA with SILVA taxonomy assignments (6.3% ± 4.5% SD) (**Fig. 2E**). This highlights the substantial impact reference database selection can have on microbiota analyses. Only reads assigned a named species from the specified taxonomy database (Greengenes or SILVA for SRA, Athena for LRA) were considered to have species-level taxonomic assignments. LRA analysis, unlike SRA analysis, could provide strain-level resolution based on the added coverage of the 2.5 kb 16-ITS-23S region. For the 12 samples analyzed with LRA, 13.4% (± 7.9% SD) of reads were assigned a specific strain designation (**Fig. 2E**).

Both metagenomic approaches assigned reads to species-level taxonomy. SRM reads were assigned NCBI taxonomy with Kraken2 and Bracken, and LRM reads were assigned taxonomy with MEGAN-LR using GTDB (**Methods**). Like 16S rRNA amplicon sequencing, metagenomics sequencing can characterize Archaea. For example, the archaea species *Methanobrevibacter smithii* was detected with SRM and LRM in both samples from Participant 5. For LRM, this archaea species was not detected using taxonomic profiling with GTDB, but it was identified through taxonomic profiling with the NCBI database, and it was also recovered through genome assembly and binning.

### Consistent taxonomic changes across methods

The clinical trial participants evaluated for this comparison displayed detectable changes in their gut microbiota from baseline to the end of STMC-103H treatment. For example, the relative abundance of genera (from LRM with GTDB taxonomy assignments) changed in Participant 3; for example, the abundance of *Agathobacter* increased while the abundance of *Anaerobutyricum* decreased (**Fig. 2F**). For each sequencing method, differential abundance analysis was performed to determine whether specific bacterial families changed in abundance from baseline across individuals (**Methods**). The most differentially abundant families with p-values less than 0.2 for SRA, LRA, and SRM, and p-values less than 0.25 for LRM are shown in **Table S2**. Eight families increased or decreased from baseline to treatment across two or three methods. Of these eight families, six showed the same directionality across methods, including increases in Akkermansiaceae, Bacteroidaceae, Oscillospiraceae, Christensenellaceae, and Clostridiales Family XIII, and decreases in Streptococcaceae (**Fig. 2G**).

### Detection of active ingredient bacterial strain in clinical fecal samples

An essential component of LBP development is the ability to track active ingredient strains after administration. While qPCR is commonly applied to detect and quantify strains in microbial diagnostics (58), microbiome sequencing can also provide insight into the fate of active ingredient strains. To confidently detect any target bacterial strain, it must be distinguished from closely related strains that may be present in a sample. The SRA ASV for the active ingredient strain DSM 33213 from the LBP STMC-103H was detected in 5 out of 6 participants after 4 weeks of treatment with STMC-103H, but not in any pretreatment baseline samples (**Fig. 3A**), suggesting that the 408 bp ASV identified in the samples originated from the LBP intervention. LRA analysis of DSM 33213 revealed it contains a single copy of the 16S-ITS-23S region with one mutation in the ITS region that enables separation of this strain from 29 other genomes of the same species in NCBI **(Fig. S2)**. DSM 33213’s 16S rRNA gene is identical to the 16S rRNA genes of 9 other NCBI genomes from the same species, so it would be impossible to distinguish from related strains based 16S rRNA sequencing alone. The active ingredient strain’s unique 16S-ITS-23S ASV was detected in all treatment sample replicates from Participant 4, but not at baseline (**Fig. 3A**). The LRA ASV for DSM 33213 could not be confidently detected in other treatment samples, potentially due to the lower sequencing depth obtained for this approach.

**Fig. 3.**
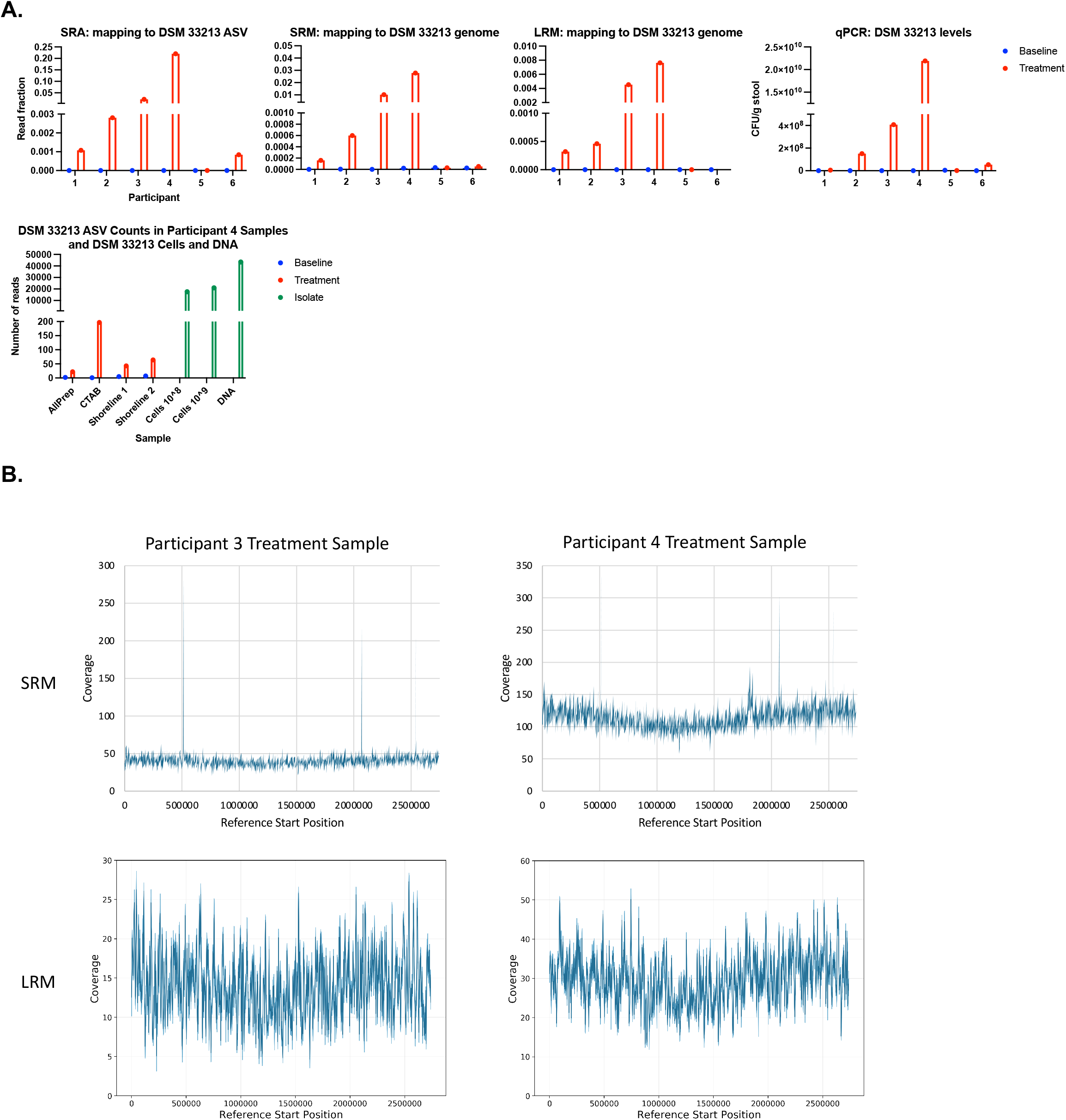
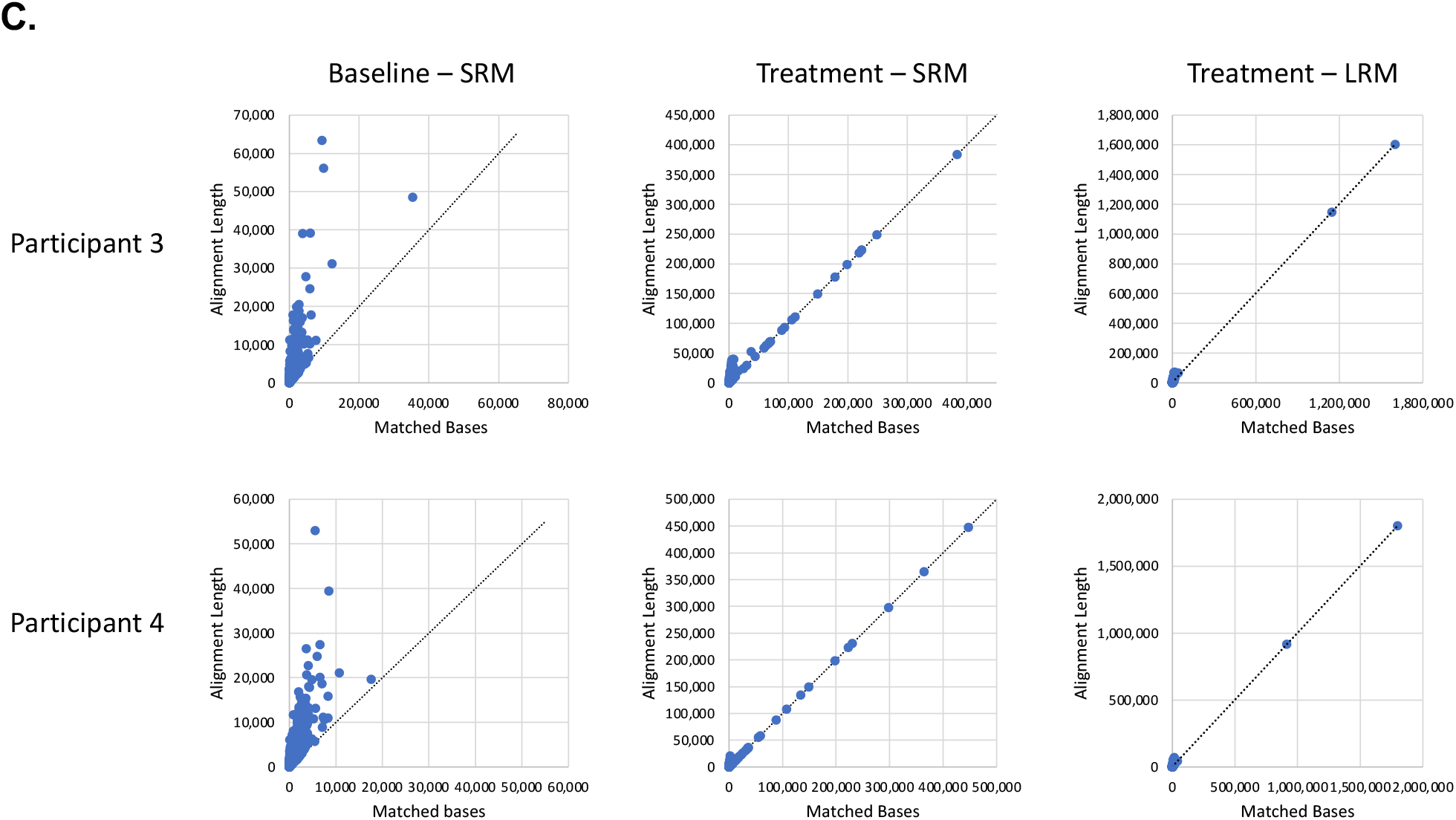
Detecting an active ingredient strain in treated samples. **(A)** The fraction of SRA reads in each sample at baseline or during treatment assigned to the 408 bp DSM 33213 ASV (top row, left graph), or the fraction of SRM and LRM reads mapping to DSM 33213’s genome (top row, center two graphs), compared to orthogonally verified levels of DSM 33213 in samples based on strain-specific qPCR (top right). The bottom graph shows levels of DSM 33213’s LRA ASV in Participant 4’s samples at baseline and treatment. All four replicates of the sample, with DNA extracted using different methods, show DSM 33213’s ASV present in treatment but not baseline replicates. DSM 33213 cells and DNA were included in the library preparation and sequencing as positive controls and are shown in green. DSM 33213’s ASV was not detected in other participants’ treatment samples. **(B)** Genomic coverage plots of SRM (top) and LRM (bottom) reads from Participant 3 and 4’s treatment samples. **(C)** Contigs assembled from SRM and LRM baseline and treatment samples from Participants 3 and 4 mapped to DSM 33213’s genome. Plots show the number of matched bases versus the total length of the alignment between each contig and the DSM 33213 genome. Dotted lines have a slope of 1 to show the reference for a perfect alignment.

By mapping metagenomic reads to DSM 33213’s genome, DSM 33213 was detected in most samples from treated individuals (5 of 6 samples by SRM and 4 of 5 samples by LRM) (**Fig. 3A**). Abundance trends for this active ingredient strain were consistent among analysis methods, and sequencing results were consistent with trends observed with strain-specific qPCR (**Fig. 3A**). Levels of DSM 33213 based on SRA and SRM significantly correlated with levels based on qPCR (Pearson r = 0.997, 0.94 and p < 0.0001, 0.01, respectively), and there was a trend towards a significant correlation between levels based on LRM and qPCR (Pearson r = 0.84, p = 0.07).

Interestingly, in the two samples with the highest quantification of DSM 33213 (treatment samples from Participants 3 and 4), mapping of LRM reads to DSM 33213’s genome generated a coverage pattern suggesting active replication, with a trough and a peak coverage profile corresponding to the bi-directional origin of replication (59) (**Fig. 3B**). This pattern of coverage was also clear for DSM 33213-mapped short metagenomic reads from Participant 4’s treatment sample. The trough and peak pattern was less clear for Participant 3’s lower coverage sample. In addition to metagenomic read coverage showing active replication of DSM 33213 in these two participants, assembled contigs from these two treated samples aligned with DSM 33213’s genome (**Fig. 3C**). Notably, the baseline samples from these two participants did not have any significant alignments to DSM 33213. From the two treatment samples, the highest quality match from the SRM contigs was a 0.45 Mbp contig with nearly 100% of bases matching DSM 33213’s genome, and the best match from the LRM contigs was a 1.6 Mbp alignment that mapped to DSM 33213 with 99.9% of bases matched. The top two LRM alignments belonged to a single 2.75 Mbp contig, which was itself a 98% complete metagenome-assembled genome (MAG) assigned to the same species as DSM 33213. This demonstrates that both SRM and LRM can recover long contigs (and for LRM, near-complete genomes) from non-endogenous microbes introduced into complex microbiota, with higher mapping confidence achieved with LRM assemblies.

### Functional microbiome profiles across methods

Beyond taxonomic annotation, metagenomic sequencing assesses the functional capabilities of a microbiome. This can provide insight into the mechanism of a microbiome’s impact on host health or the mechanism of action for a microbiome-targeted therapeutic. Although tools have been developed to predict microbial functional genes based on taxonomic profiles, their accuracy and utility are limited (60). These tools depend on databases with known sequenced microbial genomes, and dramatic differences have been observed in the genomic functions encoded between strains of the same species. Therefore, metagenomic sequencing remains an essential tool for directly evaluating the functional capacity of microbiome samples. We compared results from two metagenomic sequencing pipelines optimized for either short reads or long reads (**Methods**). Short and long reads have inherent differences and associated bioinformatic challenges. In this analysis, short reads were 150 bp while long reads were up to 10 kb, short reads are paired while long reads are singletons, and short reads and long reads have different error profiles. For SRM, read-based functional annotation was performed with HUMAnN2, which provides gene family abundances using UniProt 90 and pathway abundances with MetaCyc (38). For LRM, reads were aligned to the NCBI nr protein database and analyzed using MEGAN-LR (43). MEGAN-LR provides annotations from multiple databases including InterPro2GO, SEED, EC, and eggNOG (42).

To quantify the value of data from SRM and LRM pipelines, we compared the percent of samples’ reads with known functional annotations between the two pipelines. The percentage of reads with known functional annotations was substantially higher with LRM, even if the comparison was limited to annotations from one database (SEED for LRM). For all samples, an average of 34.2% (± 1.4% SD) of reads had known functional annotations with SRM, while an average of 63.2% (± 2.4% SD) of reads had known functional annotations for LRM (**Fig. 4A**). When all MEGAN-LR databases were included, 86.6% (± 3.2% SD) of long reads had known functional annotations, with an average of 2 to 4 functional annotations per read, which likely represent complete genes. To minimize differences between annotation pipelines and databases, the short metagenomic reads were assembled into contigs and analyzed using the LRM pipeline (**Methods**). An average of 43.6% (± 3.7% SD) of contigs displayed functional annotations across all databases, suggesting that the lower percentage of functional hits from SRM data is likely due to the limitations of short reads and not the annotation pipeline.

**Fig. 4.**
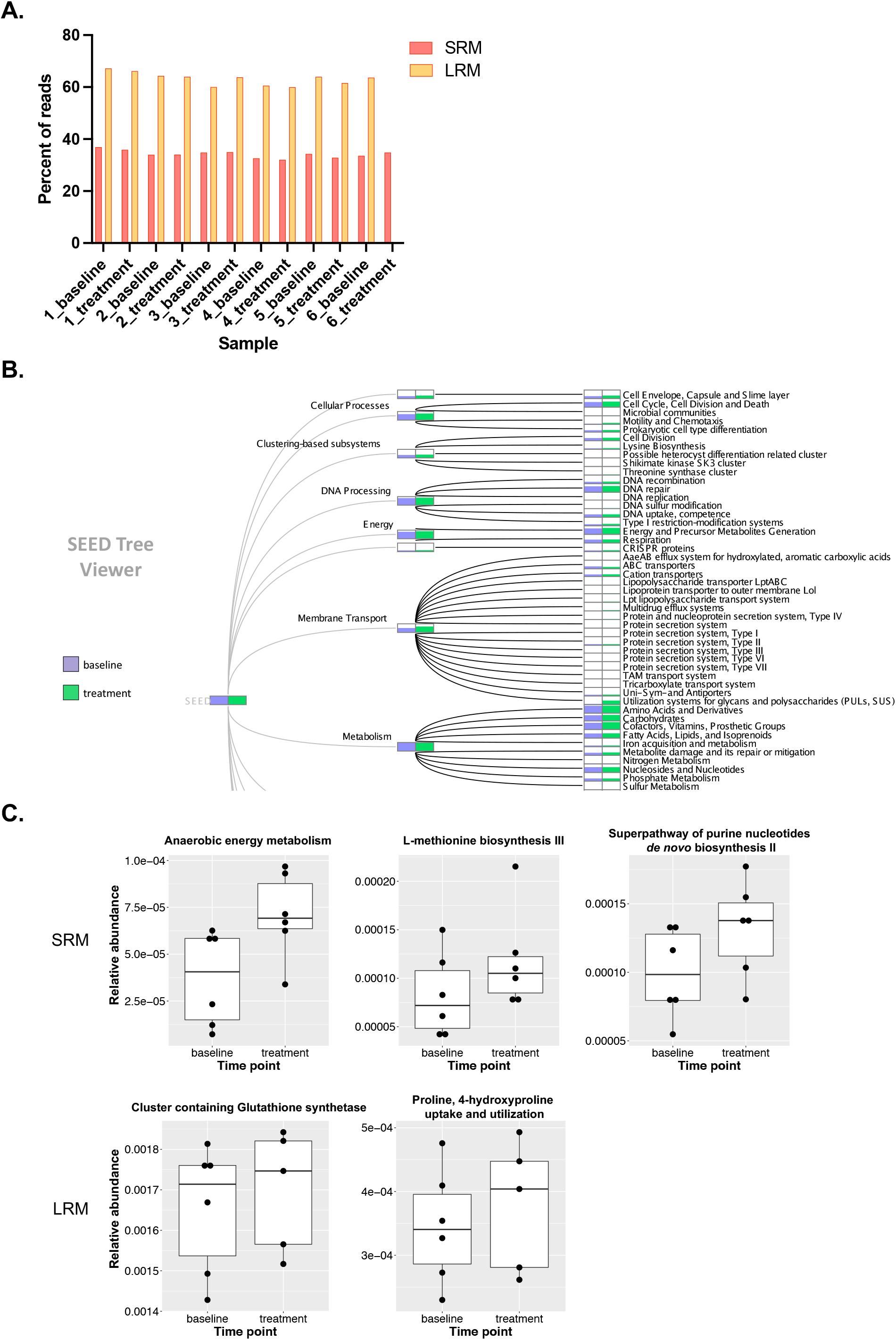
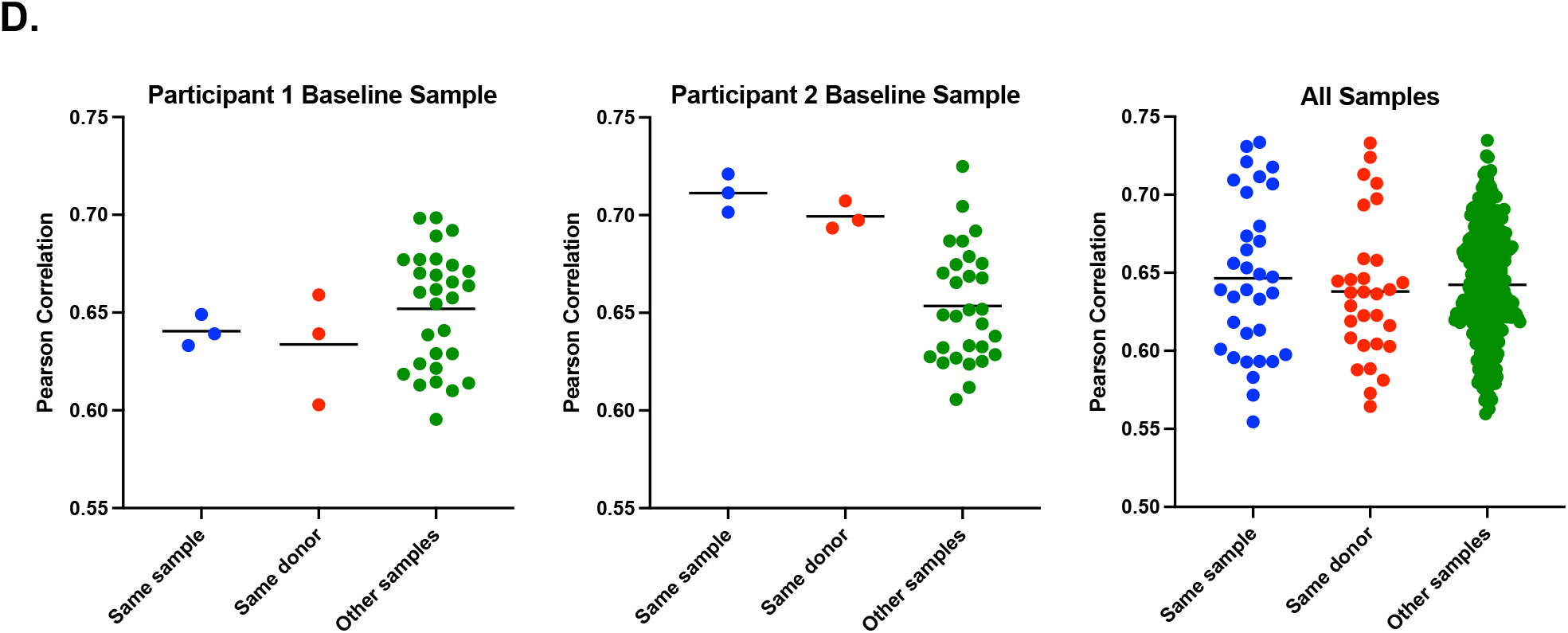
Comparison of functional annotation across methods. **(A)** The percent of reads for each sample with assigned, known functional annotation for SRM and LRM. Short reads were assigned functional annotations by HUMAnN2 using the UniProt90 database, and long reads were assigned functional annotations by MEGAN-LR using the SEED database. **(B)** Comparing the representation of SEED-based categories from LRM analysis in Participant 3’s baseline and treatment samples. **(C)** Significantly differentially abundant pathways comparing baseline and treatment samples based on SRM (top row) and LRM (bottom row). For SRM, pathways were annotated with MetaCyc, and for LRM, pathways were annotated with SEED. **(D)** Plots show the Pearson correlation between the specified sample’s LRM-based EC profile and the PICRUSt2-based EC profile based on LRA for the same sample, for the other sample from the same participant, or for all other samples. The first two plots show examples of Pearson correlations for specific samples, and the third plot shows the Pearson correlations for all samples. Horizontal lines represent the mean Pearson correlation for a given comparison.

The microbiomes of healthy individuals are relatively stable over time, but can change in response to diet, antibiotics, or other perturbations (61). This is evident in both taxonomic and functional profiles. For example, changes in the abundance of some SEED-based categories can be identified between baseline and treatment samples within a single participant (**Fig. 4B**). To determine if treatment with STMC-103H impacted the functional profiles of participants’ microbiomes in similar ways, we performed differential abundance analysis between baseline and treatment samples with SRM (HUMAnN2 pathway abundances) and LRM (SEED categories). For both SRM and LRM analyses, there were pathways that significantly increased with treatment; however, the differentially abundant pathways differed between the two methods (**Fig. 4C**).

Unlike metagenomic sequencing, marker gene-based profiling such as 16S rRNA amplicon sequencing does not provide direct information on the gene content and functional composition of a community. PICRUSt2 uses assigned taxonomy to predict the approximate functional potential of a community from marker gene sequencing profiles (34). We performed PICRUSt2 analysis with LRA data to determine if the functional profiles estimated with PICRUSt2 were similar to functional profiles determined from LRM (**Methods**). There were significant positive Pearson correlations between samples’ Enzyme Commission (EC) profiles predicted by PICRUSt2 and EC profiles from LRM. However, there were also significant positive correlations between samples from different participants, suggesting overall conserved function in similar sample types across individuals. We systematically calculated correlations between each sample’s LRM-based EC profile and the PICRUSt2-based EC profiles from A) replicates of the same sample, B) the sample from the same participant at a different time point, and C) all other samples from other participants. Surprisingly, the correlation between a sample’s LRM-based EC profile and the PICRUSt2-based profile from the same sample was not always greater than the correlation between that sample’s LRM-based EC profile and the PICRUSt2-based profiles from samples from other participants. In some cases, the mean Pearson correlation was greatest between a sample’s LRM-based EC profile and PICRUSt2-based EC profiles of samples from other participants (**Fig. 4D, left**). Overall, there was no significant difference between the Pearson correlations of samples’ LRM-based EC profile and the PICRUSt2-based profiles from the same sample, from samples from the same participant, and from samples from other participants (one-way ANOVA, p=0.67) (**Fig. 4D, right**). This suggests that, for this sample set, PICRUSt2 does not accurately estimate metagenome function.

### Diverse MAGs were recovered from metagenomic assemblies

One of the major benefits of metagenomic sequencing is the potential to recover microbial genomes that may not exist in genome databases and that provide novel information on the structure and function of the microbiome. Reads from SRM and LRM were assembled into contigs using MetaSPAdes and HiCanu, respectively. Assemblies from LRM were substantially more complete than assemblies from SRM. The average N50 for SRM assemblies was just under 7 kb, whereas the average N50 for LRM assemblies was 146 kb (**Table S3**). There was an average of 70,000 contigs per assembly from SRM and 5,400 contigs per assembly from LRM. The average largest contig per sample was 0.44 Mbp for SRM assemblies and 4.5 Mbp for LRM assemblies (**Fig. 5A**). LRM assembly generated an average of over 22 circular contigs per sample, which represent complete bacterial genomes and plasmids (**Table S3**). There were no circular contigs generated from SRM assembly. Even though SRM assemblies were significantly more fragmented, SRM and LRM total assembly lengths were comparable for each sample (**Fig. 5A**). The two exceptions, where the total assembly length was significantly shorter for SRM, were the baseline samples from Participants 2 and 6. These two samples had the fewest number of short metagenomic reads (12.4 million and 23.6 million paired reads, respectively), highlighting the need for greater sequencing depth for SRM if metagenomic assembly is desired. The average GC content of SRM and LRM assemblies were similar (45.9% and 46.1%, respectively), suggesting that there were no drastic compositional differences between the two datasets.

**Fig. 5.**
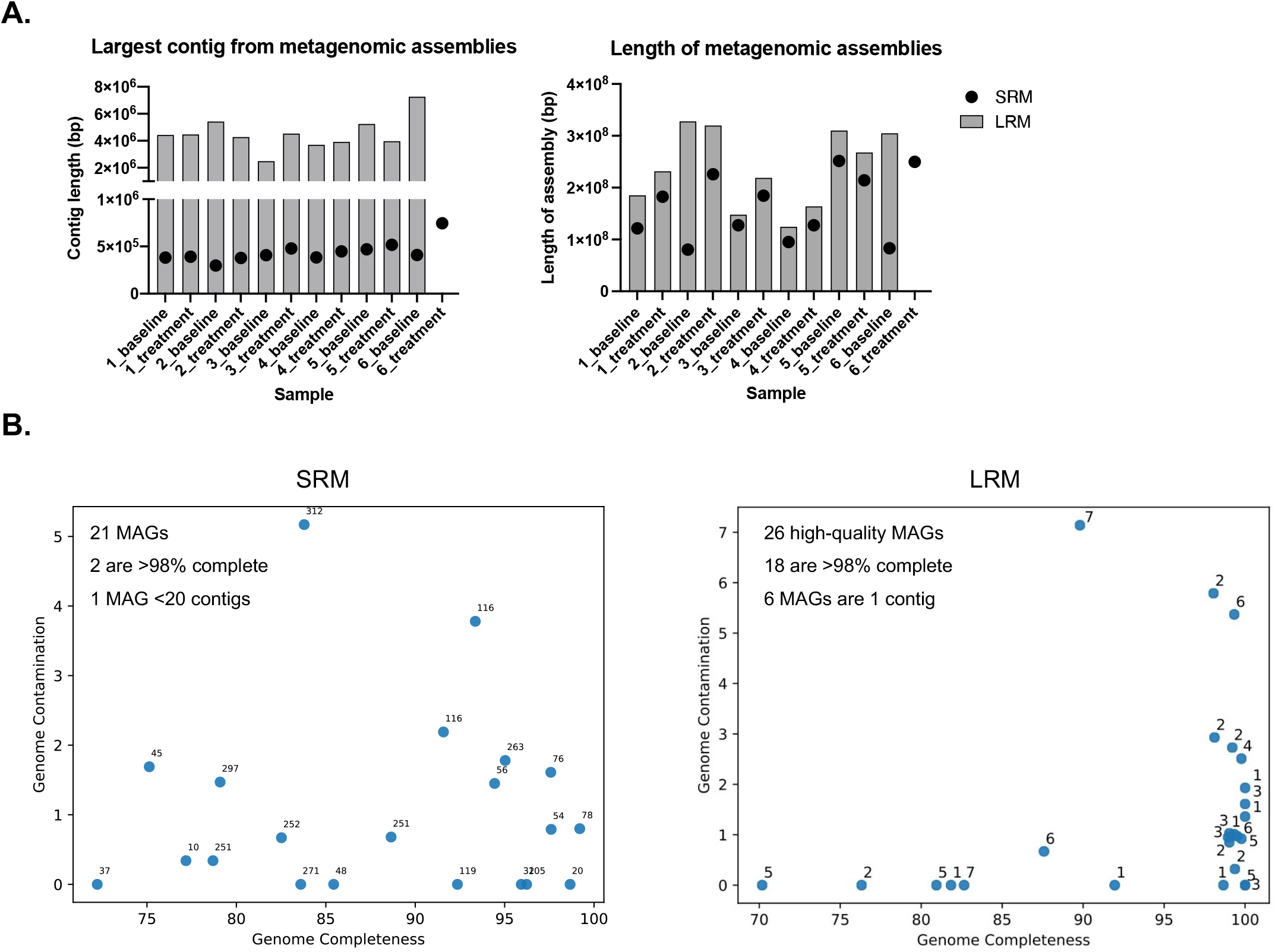
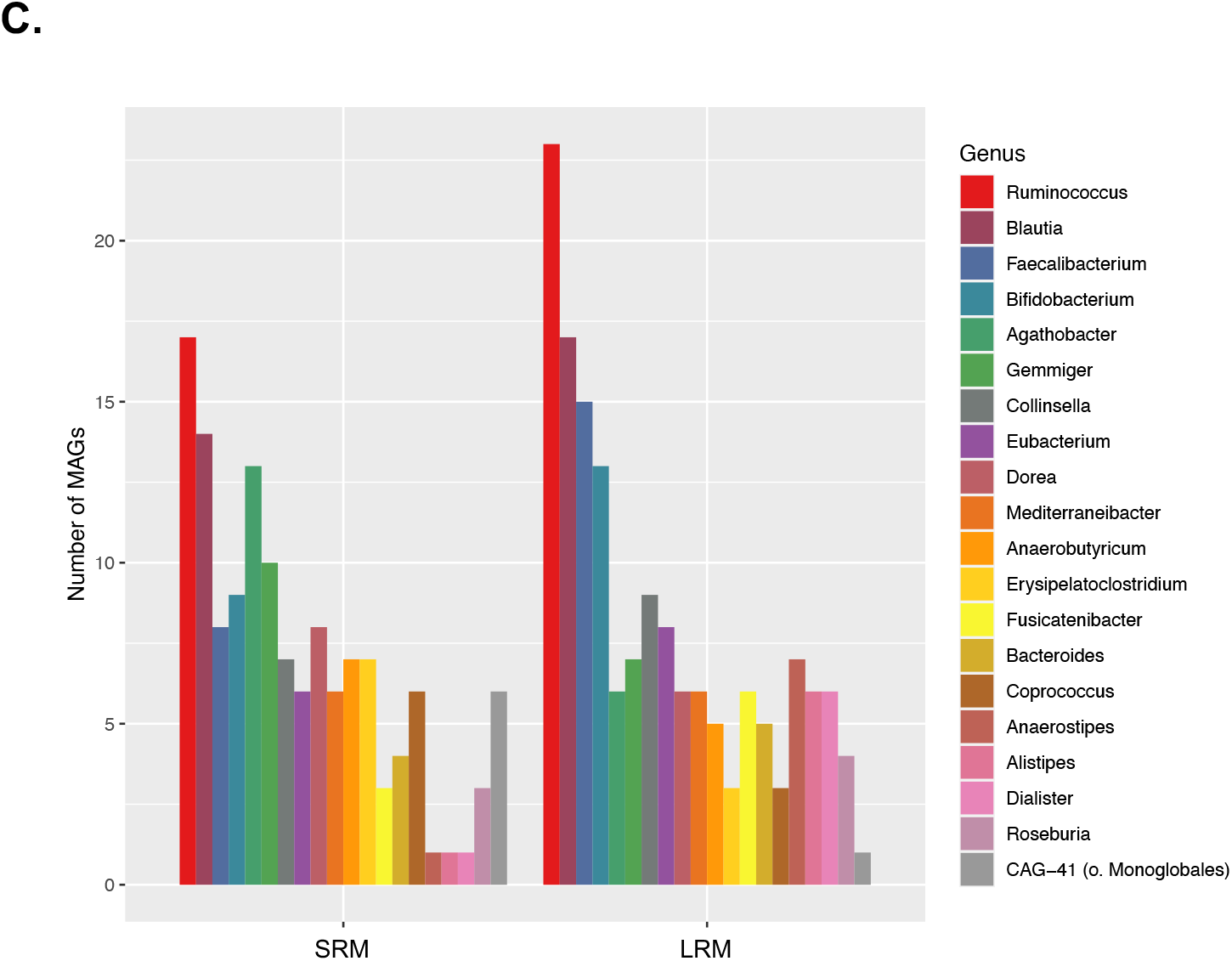
Comparison of metagenomic assemblies and metagenome assembled genomes (MAGs) for SRM and LRM. **(A)** Length of the longest contig in bp from each sample’s metagenomic assemblies from SRM and LRM (left). Total length of each sample’s metagenomic assemblies for SRM and LRM (right). **(B)** MAGs passing filtering criteria from Participant 5’s baseline sample from SRM (left) and LRM (right). **(C)** The number of MAGs passing filtering from all samples for the 20 genera with the greatest number of recovered MAGs, for SRM (left) and LRM (right).

For both SRM and LRM, genome binning and evaluation was performed using PacBio’s HiFi-MAG-Pipeline (**Methods**). Genome bins were filtered to generate high-quality metagenome assembled genomes (MAGs). The filtering criteria were relaxed for SRM to accept MAGs with 500 or fewer contigs, instead of 10 contigs or fewer as specified in the original LRM pipeline. There were no SRM bins with fewer than 10 contigs. Using the relaxed filtering criteria for SRM, each sample had an average of 17 MAGs for SRM and 18 high-quality MAGs for LRM. MAGs from LRM were substantially more complete and less fragmented than MAGs from SRM. For Participant 5’s baseline sample, there were 21 and 26 MAGs passing filtering from SRM and LRM, respectively (**Fig. 5B**). For this sample, there were only two SRM MAGs that were more than 98% complete, while there were 18 LRM MAGs that were more than 98% complete. Moreover, 6 LRM MAGs consisted of single contigs. There was only one SRM MAG with fewer than 20 contigs, while all the LRM high-quality MAGs by definition had fewer than 10 contigs.

The MAGs recovered from SRM and LRM encompassed a diverse range of taxa, and there was a high degree of overlap between methods. For example, for Participant 5’s baseline sample, there were 13 MAGs across both sequencing methods assigned the same species-level GTDB taxonomy (**Table S4**). There were 62 and 53 unique genera recovered from all samples with SRM and LRM, respectively. The five genera with the highest number of MAGs across both methods were *Ruminococcus* (40 MAGs), *Blautia* (31), *Faecalibacterium* (23), *Bifidobacterium* (22), and *Agathobacter* (19). **Figure 5C** shows the top 20 genera with the greatest number of MAGs recovered from both methods. LRM recovered a greater number of *Ruminococcus, Blautia, Faecalibacterium* and *Bifidobacterium* MAGs, while SRM recovered a greater number of *Agathobacter* and *Gemmiger* MAGs. Overall, both methods recovered similar numbers and comparable diversity of MAGs, although the MAGs were significantly more complete with LRM.

## Discussion

Taxonomic and functional profiling studies of the human microbiome largely rely on short-read DNA sequencing, despite its shortcomings. Recent improvements in long-read sequencing technology provide a promising, but more costly, alternative to traditional short-read sequencing that could potentially change the microbiome profiling landscape. To our knowledge there are no studies applying HiFi long-read sequencing to clinical microbiome samples, and this comparative evaluation illustrates the strengths and weaknesses of short-read and long-read approaches for clinical microbiome analysis and highlights the importance of choosing an approach appropriate for the research question. For taxonomic profiles—the most sought microbiome information—any of the four methods applied in this comparison can be used, since the overall proportions of phyla and families were similar across methods. The most apparent difference was LRA generating a greater proportion of unknown phyla due to the high proportion of reads annotated as bacterium LF-3. This human gut isolate is not classified at higher taxonomic levels on NCBI (Accession No. PRJEB6481), but in GTDB, it is classified as the species *Faecalibacillus intestinali*, in the phylum Firmicutes and family Erysipelatoclostridiaceae. If LRA had used GTDB taxonomy for bacterium LF-3, the overall taxonomic profiles of the samples for LRA would more closely resemble other methods at the phylum level, but not at the family level (**Fig. 2A**). Differences in taxonomic profiles across methods could be due to a variety of factors aside from database differences, including DNA extraction methods, amplification bias, and downstream analysis. However, all four methods preserved relationships between the samples.

For microbiota taxonomic profiles, taxonomic resolution is an important factor to consider when choosing a sequencing method, as increasing evidence shows that strains are the functional units of the microbiome (62). For example, although the species *Cutibacterium acnes* is a dominant skin commensal, a study showed based on full-length 16S rRNA sequencing and metagenomics, that the distribution of *C. acnes* strains significantly differed between acne patients and healthy individuals (63). Metagenomic sequencing typically provides greater taxonomic resolution than 16S rRNA sequencing, and in our analysis both LRM and SRM generated species-level taxonomic profiling. For SRA, the fraction of reads assigned to species was considerably higher when the Greengenes taxonomy database was used instead of the SILVA taxonomy database. This highlights the significant impact of the chosen reference database. LRA, unlike all other methods, provided strain-level resolution for an average of 13.4% of reads across all samples. In addition, LRA provided the greatest resolution in identifying the highest number of unique taxa per sample.

Tracking active ingredient strains is essential for measuring their impact on the native host microbiome and their fate in the gut (e.g., engraftment or clearance). All four microbiome analysis methods applied here could detect an LBP active ingredient strain in treated samples, and relative levels of this strain in treated samples were comparable across methods. Presence of the active ingredient strain was orthogonally verified with strain-specific qPCR. LRA analysis showed that the active ingredient strain DSM 33213’s ASV contains a single mutation that enables its distinction from all other known genomes of the same species. If there were a closely related strain present in the microbiota, DSM 33213 could not be confidently detected with SRA alone since the full-length 16S rRNA gene is identical to 9 other strains. Although LRA had high sensitivity in detecting DSM 33213, its specificity was lower since DSM 33213 was not detected in most treated samples. For high strain sensitivity and specificity with LRA, higher read coverage is likely necessary. Overall, LRA is a cost-effective method for tracking closely related strains in microbiota samples without the need for costly metagenomic sequencing. The StrainID method has been successfully applied to tracking closely related strains in the fecal microbiomes of premature infants (20). Importantly, STMC-103H is an LBP containing multiple strains. Only DSM 33213 could be reliably detected with sequencing methods. Other STMC-103H strains were detected in treatment samples with qPCR, but not with sequencing. Due to its higher sensitivity, qPCR should remain the gold standard for detecting specific strains. Sequencing can likely only detect strains above a certain threshold of abundance in the gut. However, sequencing can provide additional information on detected higher abundance active ingredient strains.

Targeted culturing is the primary method applied to understand the true viability, and not just presence of DNA, of active ingredient strains after administration. Here we show that metagenomics can provide information on the replication activity of an active ingredient strain from stool DNA. SRM and LRM reads mapping to DSM 33213’s genome generated peak-and-trough coverage patterns, indicative of an actively replicating strain. The genomic location of the coverage trough also matched between SRM and LRM. This provides evidence that this active ingredient strain can actively replicate in the host intestine. If the same coverage pattern of DSM 33213 persisted after cessation of LBP treatment, this would provide evidence for “engraftment” of the active ingredient strain. In a subset of participants in the trial, levels of DSM 33213 measured by strain-specific qPCR remained high one month after cessation of treatment, providing evidence for engraftment. Mapping SRM reads to DSM 33213’s genome produced spurious spikes in coverage, likely representing areas of the genome highly conserved among more distantly related organisms. The absence of these spikes with LRM mapping highlights the increased confidence inherent in mapping long reads. Assembling reads into contigs further increases mapping confidence; both SRM and LRM resulted in high-confidence matches between contigs and active ingredient strain genomes in treated samples, but not in baseline pretreatment samples. The longest alignments for SRM contigs were less than 500 kb, while the longest alignments for LRM contigs were over 1.6 Mbp, increasing confidence in active ingredient strain detection with the LRM.

Since metagenomic sequencing provides direct information about microbial gene content, it can help elucidate the mechanism of the microbiome’s contribution to health or disease or the potential mechanism of action of an LBP. For example, metagenomics analysis of microbiome samples from patients with Behcet’s disease (a rare disorder that causes blood vessel inflammation and skin sores and lesions) compared to healthy controls revealed enrichment of *Bilophila* species and opportunistic pathogens and increased microbial functions including oxidation-reduction and type III and IV secretion systems in patients with the disease (64). High-confidence functional annotations depend on high-quality sequence data. Comparing functional microbiome profiles across methods, the percentage of reads with known functional annotations was substantially higher with LRM (63% vs. 34%). Even though the cost of LRM is higher, the amount of useful data is substantially greater and requires less bioinformatic processing since HiFi reads are high-quality consensus reads. Long reads contain multiple genes in their original genomic context, increasing confidence in their taxonomic and functional annotation. There was no substantial overlap in differentially abundant pathways across methods, which could be attributed to the limited number of samples used in this analysis, differences in reference databases, and the smaller fraction of data with known annotations for SRM. PICRUSt2-based EC profiles from LRA data were not more similar to the LRM-based EC profiles from the same sample than to EC profiles from different samples. This highlights the limitation of taxonomy-based functional estimation and the importance of metagenomics for functional inference of the microbiome.

Metagenomic assemblies from SRM and LRM generated comparable assembly lengths for each sample, showing that the total unique genomic content was captured with both methods. However, LRM assemblies were substantially more complete than SRM assemblies. With relaxed filtering criteria for SRM MAGs, both methods produced comparable numbers of MAGs with overlapping taxonomy. However, only LRM binning produced high-quality MAGs consisting of single contigs, representing near-complete genomes extracted from complex microbial communities.

A notable challenge in comparing sequencing methods is that each method requires analysis pipelines and databases that are optimized for the distinct nature of the raw sequencing data. The impact of chosen databases was especially obvious when different databases were applied to the same sequencing method. For example, with SRA sequencing, using the Greengenes database increased the fraction of reads assigned to species compared to the SILVA database. Also, within LRM, *Methanobrevibacter smithii* was not identified when GTDB was used for taxonomic annotation but was identified when the NCBI taxonomic database was applied. For functional annotation of metagenomic data, independent of the analysis method, the proportion of data with known functional annotations was substantially higher with LRM than SRM. To overcome pipeline differences for this analysis, contigs assembled from SRM were analyzed using the LRM functional profiling pipeline. Even when the same pipeline was used, the proportion of assembled SRM contigs with functional annotations was still lower than the proportion of LRM reads with functional annotations.

Microbiome researchers and LBP developers must weigh the pros and cons of microbiome analysis methods to answer their core research questions. To our knowledge, this is the first comprehensive comparison of long-read and short-read approaches for microbiome characterization using samples from a live biotherapeutic intervention study. As sequencing technologies evolve, it is increasingly clear that SRA sequencing falls short of providing the high-resolution or functional microbiome information necessary to inform the development of microbiome-based interventions. Although the application of LRA sequencing provides taxonomic profiles at strain-level resolution, the lack of direct functional information is still a notable caveat of all amplicon-sequencing approaches. While LRM generated the highest amount of functionally annotated data in this study, SRM provided similar, though less complete, microbial genome recovery. While all approaches applied here could detect similar trends in the abundance of a target LBP strain, LRM sequencing provided the highest confidence in identifying an active ingredient strain, including providing evidence of active replication.

## Abbreviations

SRA: short-read amplicon sequencing
LRA: long-read amplicon sequencing
SRM: short-read metagenomic sequencing
LRM: long-read metagenomic sequencing
LBP: live biotherapeutic product
CCS: circular consensus sequencing
HiFi: high fidelity
NGS: next-generation sequencing
MAG: metagenome-assembled genome

## Author contributions

J.L.G.: Conceptualization, Data curation, Formal analysis, Investigation, Writing-original draft, Visualization. D.M.P.: Data curation, Formal analysis, Investigation, Software, Visualization, Writing-review and editing. M.D.D.: Project administration, Formal analysis, Visualization, Writing-review and editing. E.J.: Investigation, Formal analysis, Writing-review and editing. S.C.: Investigation. D.G.: Investigation, Writing-review and editing. M.A.: Project administration, Writing-review and editing. R.V.: Conceptualization, Supervision, Writing-review and editing.

## Conflicts of interest

J.G. and R.V. are employees and shareholders of Siolta Therapeutics, a clinical stage biotech company developing LBPs for the prevention and treatment of inflammatory diseases. M.D.D. is a founder and shareholder of Shoreline Biome, a biotech company developing products to advance microbiome research. E.J. and D.G. are employees and shareholders of Shoreline Biome. D.M.P., S.C., and M.A. are employees and shareholders of PacBio. All authors are committed to making the data and methods publicly available.

## Funding information

This work received no specific grant from any funding agency.

## Ethical approval

Stool samples were obtained after Institutional Review Board approval of the clinical trial protocol. Informed consent was collected from all clinical trial participants.

## Notes

https://github.com/jeanette-gehrig/Gehrig_et_al_sequencing_comparison

