## Supplemental figures and tables for "Finding the right fit: A comprehensive evaluation of short-read and long-read sequencing approaches to maximize the utility of clinical microbiome data"

### Supplementary Figures and Tables

**Fig. S1.** Scatterplots for all four sequencing methods show the number of reads sequenced versus the number of unique taxa detected for each sample. None of the Pearson correlations are significant.

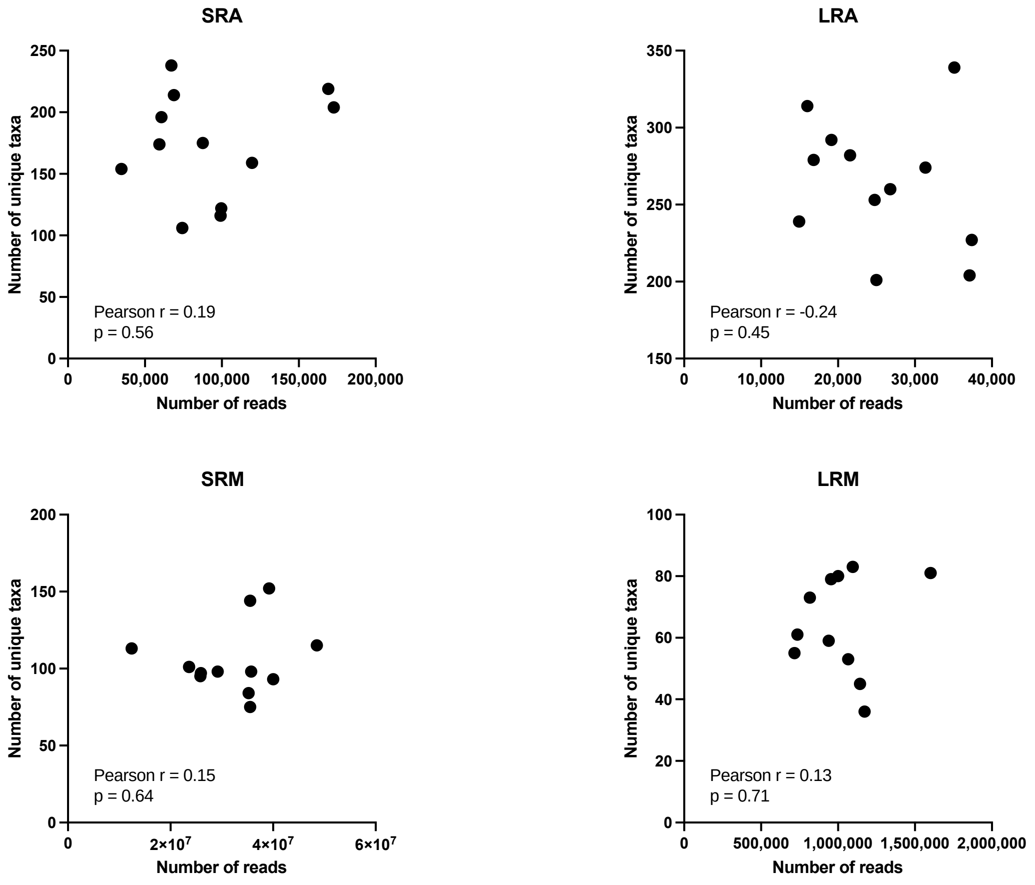


**Fig. S2. Comparison of DSM 33213's ASV to closely related genomes. (A)** LRA phylogenetic tree based on alignment of DSM 33213's ASV to the 29 most closely related genomes from NCBI. DNA alignments in Jalview were used to calculate a Neighbor Joining tree, with distances shown. DSM 33213's ASV and the most closely related sequence (Related_genome_A) from NCBI are shown in red text. **(B)** DSM 33213's 2534 bp ASV was aligned with the top hit of the 29 closely related genomes in NCBI. The deletion site in DSM 33213's ASV sequence is in a non-coding site just outside the tRNA Ile gene in the ITS region, denoted with a red box. Related Genome A was 2536 bases long and contained an insertion in the same non-coding region between the tRNA Ile and 23S gene, closer to the 23S gene. All other LRA sequences for closely related genomes were 2535 bases long and differed from DSM 33213's ASV with multiple base substitutions (not shown).

**A.**

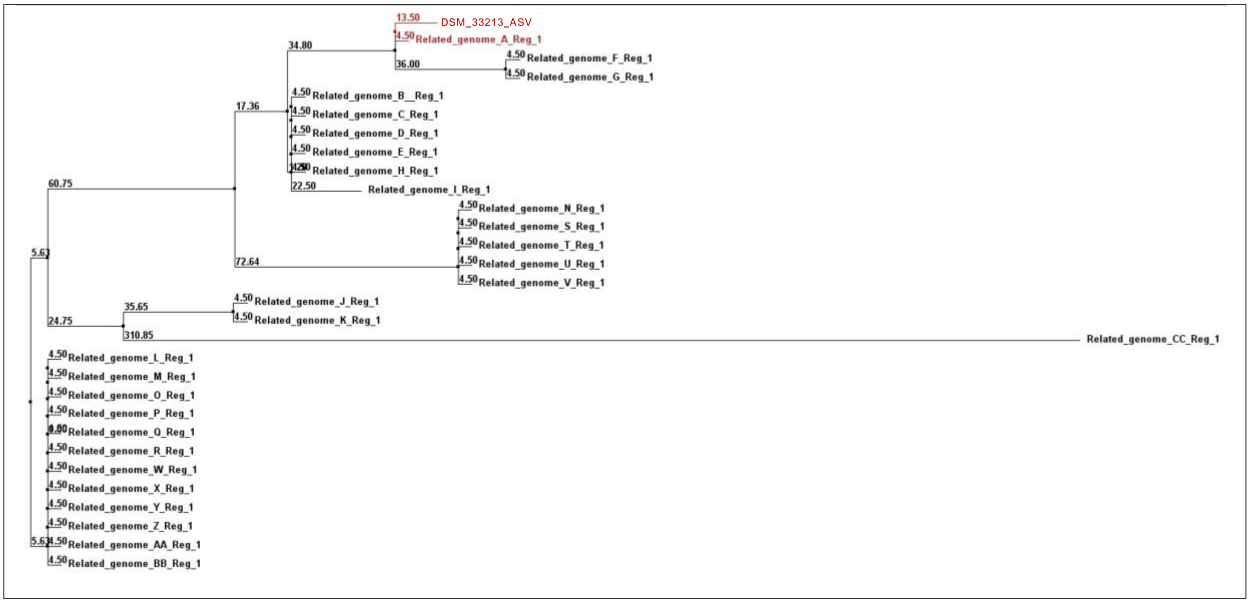

13.50 DSM_33213_ASV
4.50 Related_genome_A_Reg_1
34.80
36.00
4.50 Related_genome_F_Reg_1
4.50 Related_genome_G_Reg_1
17.36
4.50 Related_genome_B_Reg_1
4.50 Related_genome_C_Reg_1
4.50 Related_genome_D_Reg_1
4.50 Related_genome_E_Reg_1
Related_genome_H_Reg_1
22.50 Related_genome_I_Reg_1
60.75
4.50 Related_genome_N_Reg_1
4.50 Related_genome_S_Reg_1
4.50 Related_genome_T_Reg_1
4.50 Related_genome_U_Reg_1
4.50 Related_genome_V_Reg_1
72.64
5.63
35.65
4.50 Related_genome_J_Reg_1
4.50 Related_genome_K_Reg_1
24.75
310.85
Related_genome_CC_Reg_1
4.50 Related_genome_L_Reg_1
4.50 Related_genome_M_Reg_1
4.50 Related_genome_O_Reg_1
4.50 Related_genome_P_Reg_1
Related_genome_Q_Reg_1
4.50 Related_genome_R_Reg_1
4.50 Related_genome_W_Reg_1
4.50 Related_genome_X_Reg_1
4.50 Related_genome_Y_Reg_1
4.50 Related_genome_Z_Reg_1
4.50 Related_genome_AA_Reg_1
4.50 Related_genome_BB_Reg_1


**B.**

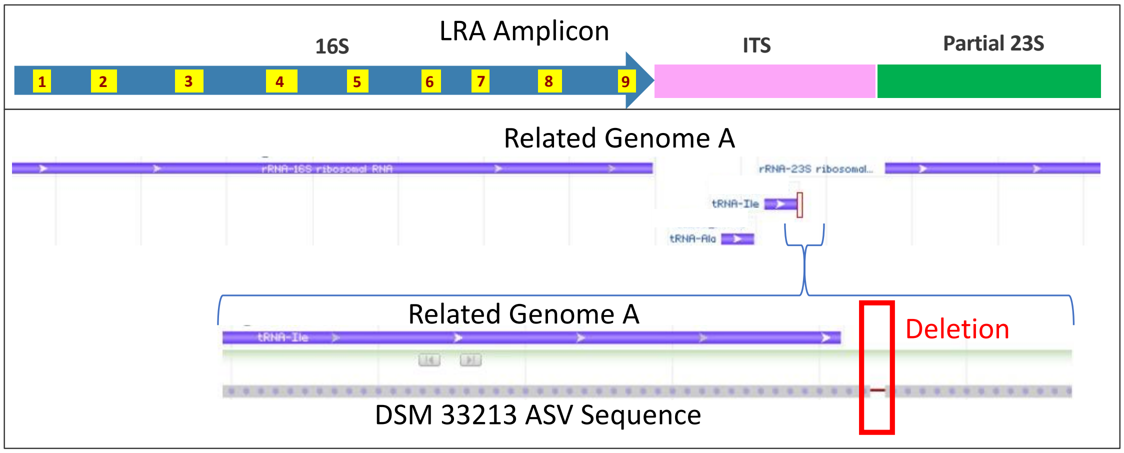

LRA Amplicon
16S
ITS
Partial 23S
1
2
3
4
5
6
7
8
9
Related Genome A
rRNA-16S ribosomal RNA
rRNA-23S ribosomal...
tRNA-Ile
tRNA-Ala
Related Genome A
tRNA-Ile
Deletion
DSM 33213 ASV Sequence

**Table S1. Summary of the DNA extraction, library preparation, data analysis methods, and databases used across the four sequencing and analysis methods.**

| Method | DNA extraction method | Library preparation | Data analysis | Taxonomy assignment databases | Functional annotation databases |
|---|---|---|---|---|---|
| SRA | CTAB(1) | Illumina protocol, Nextera adapters | QIIME 2: DADA2 plug-in to generate ASVs | GreenGenes (13_8, 99%), SILVA (132, 99%) | |
| LRA | CTAB(1), Qiagen AllPrep PowerFecal, Shoreline Complete Rapid Lysis | StrainID, SMRTbell | SBanalyzer with Athena, or DADA2 analysis to generate ASVs | Athena database (Shoreline Biome) | PICRUSt2 with KO, EC |
| SRM | CTAB(1) | Scaled-down Nextera XT | Adapters trimmed and human reads removed (BBTools), Kraken 2 and Bracken for species abundance, HUMAnN2 for functional annotation | NCBI, RefSeq | UniRef90 protein database for gene families, MetaCyc for pathways |
| LRM | Qiagen AllPrep PowerFecal | SMRTbell | Reads aligned to NCBI nr protein database with DIAMOND, taxonomy assignments and functional annotations with MEGAN6 | NCBI, GTDB | SEED, EC, InterPro2GO, eggNOG |

(1) CTAB DNA extraction protocol involves lysis with a buffer containing CTAB (cetyltrimethylammonium bromide) followed by phenol:chloroform DNA extraction with isopropanol precipitation

**Table S2. The most differentially abundant bacterial families in the six participants comparing treatment to baseline, across all four sequencing and analysis methods.** The eight highlighted families were differentially abundant across more than one sequencing and analysis method.

| Feature | Model coefficient (effect size) with reference to baseline | Model standard error | Number of samples | Number of nonzero samples | p-value | Corrected p-value (BH) | Sequencing and analysis method |
|---|---|---|---|---|---|---|---|
| Acidaminococcaceae | -0.09 | 0.06 | 11 | 5 | 0.19 | 0.86 | LRM |
| Actinomycetaceae | -0.35 | 0.16 | 12 | 12 | 0.09 | 0.72 | LRA |
| Akkermansiaceae | 0.45 | 0.26 | 12 | 12 | 0.14 | 0.72 | LRA |
| Akkermansiaceae | 1.09 | 0.62 | 12 | 10 | 0.14 | 0.7 | SRA |
| Akkermansiaceae | 0.87 | 0.49 | 12 | 12 | 0.11 | 0.87 | SRM |
| Anaerovoracaceae | 0.24 | 0.16 | 11 | 2 | 0.18 | 0.86 | LRM |
| Bacilli unclassified | 0.29 | 0.17 | 12 | 4 | 0.12 | 0.72 | LRA |
| Bacteroidaceae | 0.44 | 0.31 | 12 | 12 | 0.18 | 0.91 | SRM |
| Bacteroidaceae | 0.49 | 0.23 | 11 | 11 | 0.10 | 0.86 | LRM |
| Caldicoprobacteraceae | 0.65 | 0.29 | 12 | 8 | 0.08 | 0.72 | LRA |
| Campylobacteraceae | -0.22 | 0.12 | 12 | 12 | 0.13 | 0.87 | SRM |
| Carnobacteriaceae | -0.49 | 0.21 | 12 | 5 | 0.04 | 0.7 | SRA |
| Christensenellaceae | 0.56 | 0.23 | 12 | 11 | 0.06 | 0.72 | LRA |
| Christensenellaceae | 0.59 | 0.3 | 12 | 6 | 0.11 | 0.7 | SRA |
| Clostridiales Family XIII | 0.41 | 0.15 | 12 | 12 | 0.04 | 0.7 | SRA |
| Clostridiales Family XIII | 0.17 | 0.09 | 12 | 12 | 0.12 | 0.87 | SRM |
| Clostridiales unclassified | 0.32 | 0.17 | 11 | 9 | 0.13 | 0.86 | LRM |
| Comamonadaceae | -0.20 | 0.12 | 12 | 5 | 0.17 | 0.72 | LRA |
| Coriobacteriaceae | 0.27 | 0.15 | 12 | 9 | 0.13 | 0.72 | LRA |
| Enterobacteriaceae | -0.22 | 0.13 | 12 | 8 | 0.15 | 0.7 | SRA |
| Enterobacteriaceae | 0.07 | 0.03 | 12 | 12 | 0.04 | 0.87 | SRM |
| Eubacteriaceae | 0.26 | 0.17 | 12 | 8 | 0.18 | 0.7 | SRA |
| Gemellaceae | -0.24 | 0.11 | 12 | 5 | 0.08 | 0.7 | SRA |
| Geobacteraceae | 0.11 | 0.07 | 12 | 2 | 0.17 | 0.72 | LRA |
| Lachnospiraceae | -0.15 | 0.06 | 12 | 12 | 0.04 | 0.72 | LRA |
| Lawsonellaceae | 0.44 | 0.26 | 12 | 7 | 0.15 | 0.87 | SRM |
| Leuconostocaceae | -0.35 | 0.18 | 12 | 4 | 0.11 | 0.7 | SRA |
| Micrococcaceae | -0.43 | 0.22 | 12 | 5 | 0.08 | 0.72 | LRA |
| Micrococcaceae | 0.08 | 0.05 | 12 | 4 | 0.18 | 0.7 | SRA |
| Micrococcaceae | -0.37 | 0.18 | 12 | 12 | 0.09 | 0.87 | SRM |
| Mollicutes unclassified | 0.46 | 0.33 | 12 | 3 | 0.20 | 0.72 | LRA |
| Neisseriaceae | -0.17 | 0.12 | 12 | 2 | 0.18 | 0.72 | LRA |
| Odoribacteraceae | -0.40 | 0.26 | 12 | 7 | 0.19 | 0.72 | LRA |
| Oscillospiraceae | 0.71 | 0.39 | 12 | 11 | 0.13 | 0.72 | LRA |
| Oscillospiraceae | 0.25 | 0.13 | 12 | 12 | 0.11 | 0.87 | SRM |
| Oscillospiraceae | 0.09 | 0.06 | 11 | 5 | 0.22 | 0.86 | LRM |
| Pasteurellaceae | -0.72 | 0.46 | 12 | 6 | 0.15 | 0.7 | SRA |
| Porphyromonadaceae | 0.36 | 0.12 | 12 | 6 | 0.03 | 0.72 | LRA |
| Prevotellaceae | -0.49 | 0.2 | 12 | 6 | 0.06 | 0.7 | SRA |
| Pseudanabaenaceae | 0.27 | 0.18 | 12 | 3 | 0.19 | 0.72 | LRA |
| Pseudomonadaceae | 0.2 | 0.04 | 12 | 12 | 0.01 | 0.25 | SRM |
| Staphylococcaceae | -0.11 | 0.07 | 12 | 2 | 0.17 | 0.7 | SRA |
| Streptococcaceae | -0.38 | 0.28 | 12 | 12 | 0.2 | 0.7 | SRA |
| Streptococcaceae | -0.15 | 0.11 | 11 | 2 | 0.24 | 0.86 | LRM |
| Tissierellaceae | 0.11 | 0.07 | 12 | 2 | 0.17 | 0.72 | LRA |
| Turicibacteraceae | -0.22 | 0.12 | 11 | 3 | 0.14 | 0.86 | LRM |
| Xanthomonadaceae | -0.51 | 0.27 | 12 | 6 | 0.12 | 0.72 | LRA |

**Table S3. For each sample, summary of metagenomic assemblies from SRM and LRM.**

| | SRM - MetaSPAdes assembly | | | | | | LRM - HiCanu assembly | | | | | | |
|---|---|---|---|---|---|---|---|---|---|---|---|---|---|
| **Sample** | **Number of contigs > 500 bp** | **Largest contig** | **N50** | **Number of contigs > 50 kb** | **Total length** | **GC (%)** | **Number of contigs > 500 bp** | **Largest contig** | **N50** | **Number of contigs > 50 kb** | **Number of circular contigs** | **Total length** | **GC (%)** |
| 1_baseline | 45,087 | 382,799 | 13,429 | 334 | 121,495,177 | 46.11 | 5,135 | 4,426,676 | 100,476 | 326 | 14 | 185,210,827 | 46.5 |
| 1_treatment | 57,146 | 391,264 | 13,181 | 497 | 182,532,258 | 44.89 | 4,462 | 4,466,064 | 230,669 | 600 | 16 | 231,466,723 | 44.68 |
| 2_baseline | 60,046 | 298,123 | 1,587 | 60 | 80,455,352 | 48.46 | 7,128 | 5,425,201 | 110,354 | 1,047 | 26 | 327,828,295 | 46.14 |
| 2_treatment | 102,213 | 379,527 | 5,021 | 295 | 225,828,910 | 47.12 | 7,662 | 4,278,698 | 84,774 | 979 | 26 | 319,740,244 | 46.93 |
| 3_baseline | 50,144 | 408,049 | 8,508 | 250 | 127,346,222 | 42.97 | 4,867 | 2,482,580 | 83,017 | 338 | 24 | 147,845,990 | 45.49 |
| 3_treatment | 69,258 | 478,836 | 9,426 | 411 | 184,370,755 | 46.14 | 4,779 | 4,527,864 | 182,040 | 530 | 21 | 218,733,090 | 46.12 |
| 4_baseline | 43,617 | 384,652 | 5,644 | 218 | 95,109,160 | 44.39 | 3,345 | 3,699,694 | 91,022 | 278 | 11 | 124,622,535 | 45.85 |
| 4_treatment | 52,732 | 447,722 | 6,618 | 279 | 127,562,644 | 45.18 | 4,196 | 3,922,663 | 119,960 | 423 | 19 | 163,975,178 | 45.95 |
| 5_baseline | 112,099 | 469,277 | 5,158 | 414 | 251,479,149 | 45.79 | 6,370 | 5,247,245 | 119,615 | 812 | 29 | 310,092,326 | 45.77 |
| 5_treatment | 94,323 | 517,697 | 6,509 | 417 | 214,137,139 | 44.53 | 6,350 | 3,973,077 | 113,993 | 709 | 32 | 267,985,865 | 46.02 |
| 6_baseline | 49,532 | 409,256 | 2,824 | 78 | 83,262,378 | 49.19 | 5,091 | 7,265,491 | 369,839 | 713 | 29 | 304,712,648 | 47.71 |
| 6_treatment | 103,796 | 746,119 | 5,543 | 388 | 249,803,195 | 45.45 | | | | | | | |
| **Average** | 69,999.4 | 442,776.8 | 6,954.0 | 303.4 | 161,948,528.3 | 45.9 | 5,398.6 | 4,519,568.5 | 145,978.1 | 614.1 | 22.5 | 236,564,883.7 | 46.1 |
| **SD** | 25,649.3 | 111,412.8 | 3,644.0 | 135.9 | 63,804,606.8 | 1.7 | 1,316.6 | 1,201,491.5 | 86,632.2 | 263.5 | 6.8 | 74,119,789.2 | 0.8 |

**Table S4. Overlap of taxonomy assigned to MAGs passing filtering criteria from SRM and LRM assemblies from participant 5's baseline sample.**

| MAG taxonomy | SRM | LRM |
|---|---|---|
| s__Methanobrevibacter_A smithii | ✓ | ✓ |
| s__Bifidobacterium adolescentis | | ✓ |
| s__Bifidobacterium angulatum | | ✓ |
| s__Bifidobacterium longum | ✓ | ✓ |
| g__Collinsella | ✓ | ✓ |
| s__Prevotellamassilia sp900539625 | | ✓ |
| s__CAG-279 sp000437795 (f__Muribaculaceae) | | ✓ |
| s__Clostridium sp000435835 | ✓ | |
| s__Agathobacter faecis | ✓ | ✓ |
| s__Anaerostipes hadrus | | ✓ |
| s__Anaerobutyricum hallii | ✓ | |
| s__Bariatricus comes | ✓ | ✓ |
| s__Blautia_A sp900066205 | | ✓ |
| s__Blautia_A wexlerae | ✓ | |
| s__CAG-194 sp000432915 (f__Lachnospiraceae) | ✓ | ✓ |
| s__CAG-45 sp900066395 (f__Lachnospiraceae) | ✓ | ✓ |
| s__Fusicatenibacter saccharivorans | | ✓ |
| s__Mediterraneibacter faecis | | ✓ |
| s__Roseburia intestinalis | | ✓ |
| s__Roseburia inulinivorans | | ✓ |
| s__Coprococcus_A catus | ✓ | |
| s__Dorea longicatena | ✓ | |
| s__UMGS1375 sp900066615 (f__Lachnospiraceae) | ✓ | ✓ |
| s__CAG-177 sp003514385 (f__Acutalibacteraceae) | ✓ | ✓ |
| s__Eubacterium_R sp000436835 | ✓ | ✓ |
| s__Ruminococcus_E bromii_B | ✓ | ✓ |
| s__Faecalibacterium prausnitzii | | ✓ |
| s__Faecalibacterium prausnitzii_D | | ✓ |
| s__CAG-110 sp000434635 (f__Oscillospiraceae) | ✓ | |
| s__Gemmiger qucibialis | ✓ | |
| s__Ruminiclostridium_E siraeum | ✓ | ✓ |
| s__Ruminococcus_C sp000433635 | | ✓ |
| s__Erysipelatoclostridium sp000752095 | ✓ | |
| s__Holdemanella sp002299315 | ✓ | ✓ |
| **Total** | **21** | **26** |
| **Shared** | **13** | |
